# SydR, a redox-sensing MarR-type regulator of *Sinorhizobium meliloti*, is crucial for symbiotic infection of *Medicago truncatula* roots

**DOI:** 10.1101/2023.10.17.562213

**Authors:** Fanny Nazaret, Davoud Farajzadeh, Joffrey Mejias, Marie Pacoud, Anthony Cosi, Pierre Frendo, Geneviève Alloing, Karine Mandon

## Abstract

Rhizobia associate with legumes and induce the formation of nitrogen-fixing nodules. The regulation of bacterial redox state plays a major role in symbiosis and <u>R</u>eactive <u>O</u>xygen <u>S</u>pecies (ROS) produced by the plant are known to activate signaling pathways. However, only a few redox-sensing transcriptional regulators (TRs) have been characterized in the microsymbiont. Here, we describe SydR, a novel redox-sensing TR of *S. meliloti* that is essential for the establishment of symbiosis with *Medicago truncatula*. SydR, a MarR-type TR, represses the expression of the adjacent gene SMa2023 in growing cultures, and this repression is alleviated by NaOCl, *tert-*butyl, or H_2_O_2_ treatment. Gels shift assays strongly suggest that SydR binds to <u>TATCGCGATA</u> motif in the *sydR*-SMa2023 intergenic region in a redox-dependent manner. Furthermore, site-directed mutagenesis demonstrated that the oxidative inhibition of SydR involves the formation of an intermolecular C16-C16 disulfide bond. The inactivation of *sydR* did not alter the sensitivity of *S. meliloti* to NaOCl, *tert*-butyl, or H_2_O_2_, nor did it affect the response to oxidants of the roGFP2-Orp1 redox biosensor expressed within bacteria. However, *in planta*, Δ*sydR* mutation impaired the formation of root nodules. Microscopic observations and analyses of marker gene expression showed that the Δ*sydR* mutant is arrested at an early stage of the bacterial infection process. Altogether, these results demonstrated that SydR is a redox sensing MarR-type TR that plays a key role in the regulation of symbiosis with *M. truncatula*.

**IMPORTANCE:** The nitrogen-fixing symbiosis between rhizobia and legumes has an important ecological role in the nitrogen cycle, contributes to nitrogen enrichment of soils, and can improve plant growth in agriculture. This interaction is initiated in the rhizosphere by a molecular dialog between the two partners, resulting in plant root infection and formation of root nodules, where bacteria reduce the atmospheric nitrogen into ammonium. This symbiosis involves modifications of the bacterial redox state in response to reactive oxygen species produced by the plant partner. Here, we show that SydR, a transcriptional regulator of the *Medicago* symbiont *Sinorhizobium meliloti*, acts as a redox-responsive repressor that is crucial for the development of root nodules and contributes to the regulation of bacterial infection in *S. meliloti / Medicago truncatula* symbiotic interaction.

## INTRODUCTION

**B**acteria have to cope with environmental variations either as free-living microorganisms or during interaction with eukaryotic organisms. The success of this adaptation depends on their ability to coordinate changes in gene expression and maintain cellular homeostasis. In particular, bacteria have evolved different transcriptional regulators able to sense variations in the level of redox-active compounds, such as Reactive Oxygen Species (ROS) and Reactive Nitrogen Species (RNS), known as key signaling molecules. Thereby, redox-responsive regulators modulate the expression of target genes involved in bacterial response to changes in redox state and in ROS-RNS regulated-cell processes (1). The important role of ROS has been particularly highlighted in beneficial and pathogenic interactions between plants and microorganism (2–4). Accordingly, the production of ROS in plants and antioxidant defense in bacteria are major determinants for the outcome of the interactions.

Soil Rhizobacteria are able to reduce atmospheric nitrogen (N_2_) into ammonia in symbiosis with a wide range of legumes, enabling them to sustain plant growth in nitrogen-limited environments. Thanks to this biological nitrogen fixation (BNF), the symbiotic bacteria contribute to nearly half of nitrogen input in crop soils, giving legume plants an agronomic advantage (5). BNF occurs in a newly emerged plant organ called nodule. Both rhizobia and host plants exhibit narrow specificity, and the interaction is initiated by an exchange of molecular signals. Flavonoids released into the rhizosphere by the host plant induce bacterial secretion of lipochito-oligosaccharides, the Nodulation Factors (NFs). NFs, in turn, trigger signal transduction pathways in plants, leading to activation of central regulators and subsequent reprogramming of root cortical cells for nodule organogenesis and bacterial infection (6). In *Medicago* sp., the root hairs curl to form an infection pocket where entrapped bacteria divide. Then, the bacteria progress toward plant cortical cells, within a host tubular structure emerging from the infection pocket called infection thread (IT). Finally, the bacteria are released inside plant cells and differentiate into nitrogen-fixing bacteroids.

ROS play an important role during all steps of symbiosis development, and changes in ROS concentration have been detected from the early stage of interaction to mature nodules (7). A recent analysis using redox biosensors has established that the microsymbiont maintains a reduced state inside IT and undergoes an oxidative upshift during bacteroid differentiation (8). Moreover, several genes involved in the antioxidant defense of the microsymbiont have been known to be required for optimal symbiosis (9). The search for central redox-sensing regulators involved in the control of the legume-rhizobia symbiosis has been undertaken for several years. In bacteria, most of the redox-sensing regulators belonging to the LysR and MarR families are involved in various processes (10). Their activity is often regulated *via* oxidative post-translational modification of specific cysteines. This mechanism enables redox-sensing regulators to rapidly trigger activation or inactivation of their target promoters. In *S. meliloti*, the conserved LysR-like regulator OxyR contributes significantly to the response to H_2_O_2_ in growing cultures (11, 12). However, no symbiotic phenotype has been associated with an *oxyR* inactivated mutant. Following a large-scale mutagenesis approach, another LysR-like redox-sensing regulator, LsrB, was identified for its important role in the efficiency of *S. meliloti* / *Medicago sativa* interaction (13). An *lsrB* deletion mutant induces ineffective, early senescing nodules with abnormal bacteroid differentiation (14). Beside LysR-like regulators, members of the MarR (<u>M</u>ultiple <u>a</u>ntibiotic resistance <u>R</u>egulator) family are widely distributed in bacteria and archaea (15). The redox-responsive MarR-type regulators are generally repressors that are inactivated upon oxidation, which triggers target gene expression. Among them, the MarR/OhrR proteins respond more particularly to peroxides (10). In *S. meliloti*, OhrR regulates *ohr* shown to be expressed in symbiosis, but neither *ohrR* nor *ohr* inactivation affects symbiosis efficiency (16). Finally, while this work was in progress, Zhang *et al*, (17) published that a deletion in SMa2020, an OhrR-like encoding gene, reduced IT formation and plant growth with no reported effect on the number of nodules in *S. meliloti* / *M. sativa* interaction.

In this work, the redox-response of MarR-type candidates in *S. meliloti* was analyzed. Two of them, encoded by SMc00146 and SMa2020 genes, were found to control the expression of sodium hypochlorite (NaOCl)-inducible genes. SMa2020 encodes a transcriptional repressor containing a redox-sensitive cysteine. In contrast with the results observed during the *S. meliloti* / *M. sativa* interaction, the inactivation of SMa2020 (called SydR for <u>Sy</u>mbiosis re<u>d</u>ox <u>R</u>egulator) resulted in the early abortion of bacterial infection with a drastic decrease in nodule number during the interaction with *M. truncatula*.

## RESULTS

### Identification of redox-sensing MarR-type regulators in *S. meliloti*

The annotation of *S. meliloti* genome (*S. meliloti* 2011 genome website, https://iant.toulouse.inra.fr) allowed the identification of 10 genes encoding MarR-type proteins with one or more cysteine(s) (Table S1). Three *S. meliloti marR-type* genes have already been studied, *i.e.* SMc00098 encoding OhrR, SMc01945 encoding Cpo, and SMb21317 encoding WggR (16, 18, 19). Of the seven candidates not yet characterized, SMc00562 and SMc04052 were ruled out because their expression is barely detected in nodules according to Roux and coll. (20). Based on Clustal-O multiple alignment analysis with predicted secondary structure (Jalview), the sequences of proteins encoded by the remaining five candidates display a typical wHTH (wing Helix-Turn-Helix) DNA-binding domain in the central part of the protein and a largely helical dimerization domain (Fig. 1A; 21). The both cysteines of SMc01908 product were found in the wHTH domain, which precludes redox-sensing activity of the cysteine. Thus, further analyses were conducted with SMa2020, SMc00146, SMc00384 and SMc03824.

**FIG 1.**
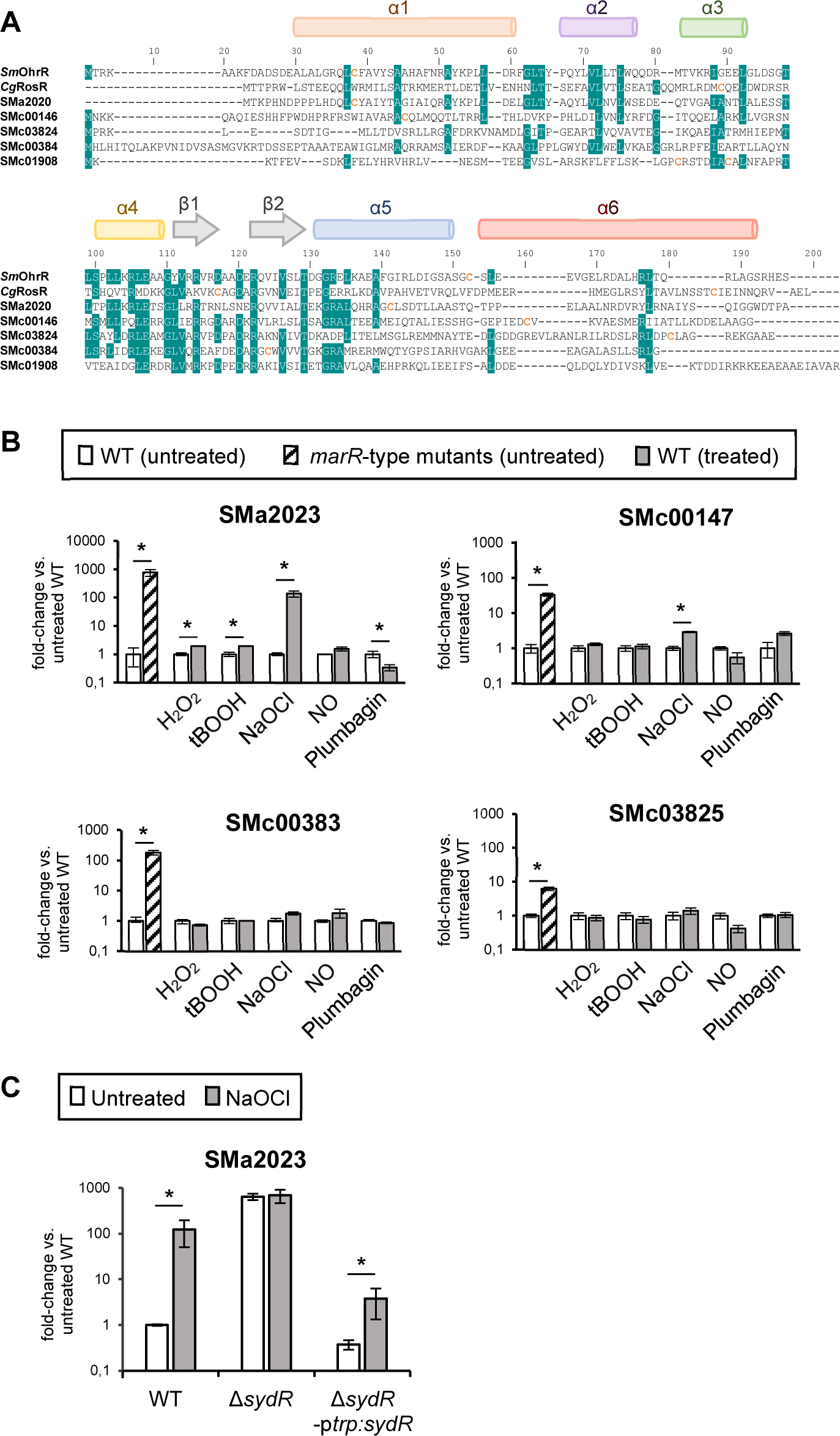
SydR is a new MarR-type redox-sensing repressor. (A) Sequence alignment of MarR-type regulators. The alignment between *S. meliloti* OhrR and SMa2020, SMc00146, SMc03824, SMc00384, SMc01908 products, and *Corynebacterium glutamicum* RosR, was generated using Clustal-W. Secondary structure elements were predicted from the sequence of SMa2020 product using Jalview (50). The HTH domain corresponds to helices α3 and α4, the wing to β-strands β1 and β2, and the dimerization domain is formed by helices α1, α5 and α6. Cysteine residues are indicated in orange. Blue shading indicates an identity ≥70% at that position. (B) Analysis of MarR-type-mediated gene expression. RT-qPCR analysis of the expression of SMa2023, SMc00147, SMc00383 and SMc03825, considered as potential target genes of SMa2020, SMc00146, SMc00384 and SMc03824 products, respectively. The expression of each gene was analyzed in WT under control condition (untreated), in the related *marR*-type derivative mutant and in WT challenged with H_2_O_2_, tBOOH, NaOCl, NO or plumbagin (treated). (C) The induction of SMa2023 expression by NaOCl is SydR-dependent. RT-qPCR analysis of SMa2023 expression in WT, Δ*sydR* (SMa2020-inactivated mutant) and Δ*sydR-*p*trp:sydR* strains under control conditions (untreated) or challenged with NaOCl. For each condition, transcription levels were normalized to those in untreated WT. The values shown are the means ± SEM of three independent experiments. Student’s *t* test was used to assess the statistical significance (*, p<0.05).

Most redox-sensing MarR are transcriptional repressors inactivated by redox-active compounds such as ROS, RNS or NaOCl. To investigate the role of the four candidates in regulating gene expression, we identified a putative target gene in close vicinity of each MarR-type encoding gene and first examined its basal expression depending on *marR*-type gene integrity. For this purpose, mutants in SMa2020, SMc00146, SMc00384 and SMc03824 were constructed. Comparative RT-qPCR experiments were performed using total RNA extracted from WT and mutant bacteria grown in M9 medium. As shown in Fig. 1B, a strong increase in specific mRNA level was obtained in the mutants as compared to the WT strain (with a fold change of 6 for SMc03825, 33 for SMc00147, 180 for SMc00383 and 773 SMa2023). These results suggest that each selected *marR-*type gene encodes a repressor that inhibits the expression of proposed target gene under basal conditions.

To test whether the repression by MarR-type proteins was dependent on redox sensing, bacterial cultures were exposed to various ROS and redox-active compounds and the effect of treatments on target gene expression was investigated. Sublethal concentrations of H_2_O_2_, *tert*-butyl hydroperoxide (tBOOH), spermine NONOate (a nitric oxide generator), NaOCl and plumbagin (an anion superoxide generator) capable of inducing gene expression were first determined, by using oxidative stress-inducible marker genes (Fig. S1). Then, the same concentrations of oxidants were applied to analyze the expression of MarR-type target genes (Fig. 1B). SMc00383 and SMc03825 expression remained unchanged regardless of the added oxidant. In contrast, the expression of SMc00147 was increased 3-fold by NaOCl treatment, and that of SMa2023 was induced 137-fold by NaOCl treatment, 2-fold by H_2_O_2_ and tBOOH, and reduced 3-fold by plumbagin addition. Therefore, the repression of SMc00147 and SMa2023 transcription can be alleviated by the addition of oxidants, supporting the hypothesis that SMc00146 and SMa2020 are redox-responsive regulators. Considering the higher induction of SMa2023 compared to SMc00147, we focused on the characterization of SMa2020, which was named SydR for <u>Sy</u>mbiosis re<u>d</u>ox <u>R</u>egulator.

To further analyze the relationship between *sydR* (SMa2020) and the NaOCl-induced expression of SMa2023, the SMa2023 expression level was determined in WT, Δ*sydR* (SMa2020 defective mutant) and in a Δ*sydR* strain expressing *sydR* under the control of the constitutive promoter p*trp* (Δ*sydR*-p*trp*:*sydR*) (Fig. 1C). A similar expression level was observed in both untreated and NaOCl-treated cultures of Δ*sydR* mutant, showing that the redox regulation of SMa2023 was lost in this strain. In contrast, the redox regulation of SMa2023 was recovered in the complemented strain Δ*sydR*-p*trp:sydR*. Altogether, these data establish that the induction of SMa2023 expression by NaOCl is controlled by SydR.

### Specific binding of SydR to the *sydR*-SMa2023 intergenic region

To test the direct regulation of SMa2023 by SydR, we analyzed the interaction between SMa2023 promoter and a His-tagged recombinant SydR purified in the presence of DTT, called SydR’ (Fig. 2). The direct binding of SydR’ to *sydR*-SMa2023 intergenic region was detected by electrophoretic mobility shift assay (EMSA) with SydR’ and a 144-bp specific DNA fragment encompassing the entire region between *sydR* and SMa2023 ORFs (Fig. 2A). The addition of increasing concentrations of SydR’ resulted in a band up-shift in a dose dependent manner (Fig. 2B). SydR’ DNA-binding was conserved in the presence of competitor poly(dI-dC) DNA and was absent with non-target DNA (internal fragment of SMa2023) demonstrating its specificity (Fig. 2C).

**FIG 2.**
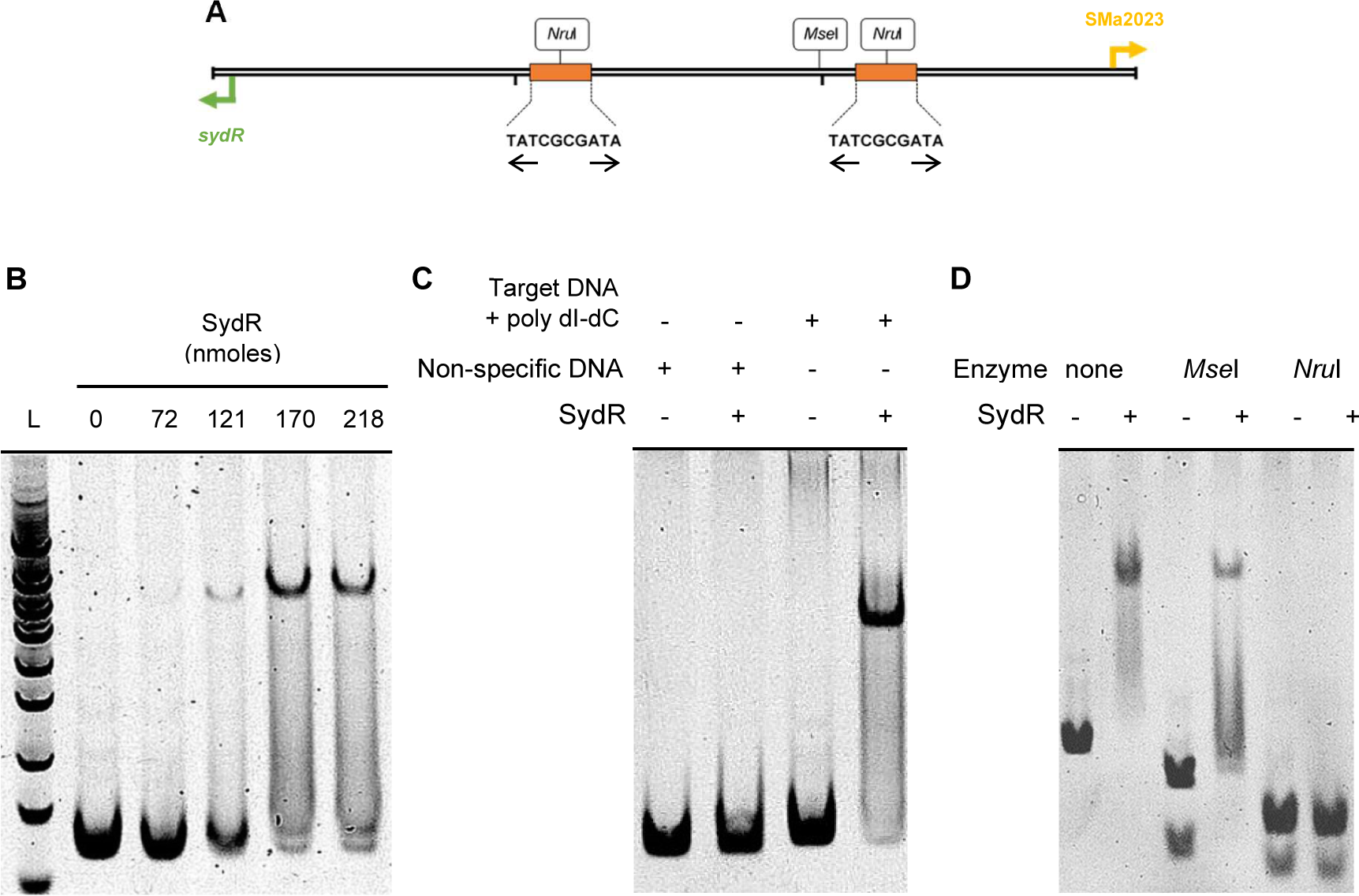
SydR binds with high specificity to the promoter region of *sydR* and SMa2023. (A) Restriction map of the 144 bp *sydR*-SMa2023 intergenic region. (B, C, D) EMSA analysis of SydR DNA-binding activity. (B) The *sydR*-SMa2023 intergenic region was incubated with increasing concentrations of SydR’, and the mixtures were separated in a polyacrylamide non-denaturing gel. (C) The *sydR*-SMa2023 intergenic region, poly (dI-dC) as competitor and non-target DNA were incubated in the presence (+) or not (-) of SydR’. (D) The *sydR*-SMa2023 intergenic region and its restriction fragments produced by *Mse*I or *Nru*I digestion were incubated with SydR’.

To identify SydR DNA-binding site(s), a search for inverted repeat motifs in the *sydR*-SMa2023 intergenic region was performed, by using an EMBOSS online tool (https://www.bioinformatics.nl/cgi-bin/emboss/palindrome). This led to the discovery of two identical 10-nucleotide sequences (<u>TATCGCGATA</u>) (Fig. 2A). A cleavage site for *Mse*I is located between the two DNA motifs, while *Nru*I cleaves each of them, creating three DNA fragments of 37 bp, 53 bp, and 54 bp respectively. Digestion of target DNA by *Mse*I prior to EMSA assay maintained the recombinant protein binding, while disruption of the inverted repeats by *Nru*I prevented upshift of resulting fragments (Fig. 2D). Overall, these data strongly suggested that SydR is capable of directly regulating SMa2023 by binding to the TATCGCGATA motif present in the *sydR*-SMa2023 intergenic region.

### Oxidative inhibition of SydR occurs via the formation of an intermolecular C16-C16 disulfide bond

The sensitivity of SydR DNA-binding activity to oxidation was analyzed *in vitro* (Fig. 3A-C). SydR’ was subjected to oxidation by H_2_O_2_, tBOOH, and NaOCl application. As shown in Fig. 3A, SydR’ bound DNA in control condition as well as after treatment with H_2_O_2_ or tBOOH (4 μM). In contrast, SydR underwent oxidative inhibition with 1 μM NaOCl. The addition of DTT (2 mM) in binding solution restored the DNA-binding activity of SydR’. These data showed that the oxidation by NaOCl impairs DNA binding in a reversible manner. Thus, the ability of SydR to bind DNA depends on the redox state of the protein, most likely via redox sensitive cysteine(s).

**FIG 3.**
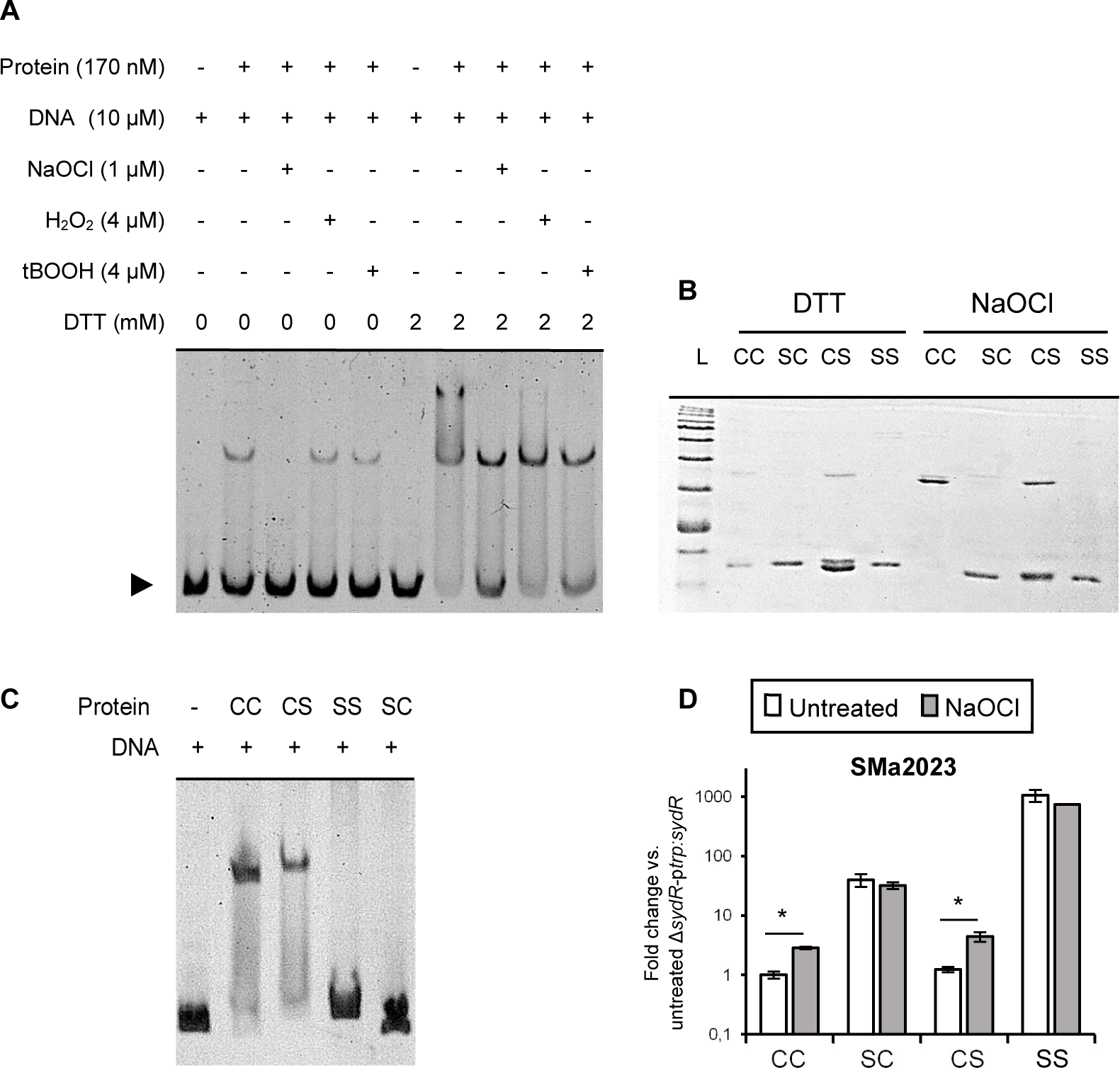
SydR binding activity is inhibited by oxidation and depends on the redox active cysteine C16. (A) EMSA with SydR’ treated or not with various oxidants, then re-reduced with DTT when indicated. Non-reducing SDS-page (B) and EMSA (C) using SydR’ WT protein (CC) and mutant variants harboring the C16S (SC), C114S (CS), and C16S&C114S (SS) mutations, treated with DTT then, when indicated, with NaOCl. (D) RT-qPCR analysis of SMa2023 expression in Δ*sydR*-p*trp:sydR* strain and Δ*sydR* derivatives encoding SydR mutant variants (SC, CS or SS). The values shown are the means ± SEM of three independent experiments. Student’s *t* test was used to assess the statistical significance of differences compared to untreated Δ*sydR*-p*trp:sydR* (*, p<0.05).

To investigate the role of cysteine residues in the structure and DNA binding activity of SydR, the two cysteine residues (C16 and C114) were replaced with serine by site-directed mutagenesis, and recombinant proteins containing one or two mutated cysteines were produced. The dimerization state of SydR was assessed by non-reducing SDS-PAGE using WT and mutant proteins purified in the presence of DTT (Fig. 3B). SydR’ contains both cysteine residues in a reduced form, resulting in a monomer with a molecular mass of 20.4 kDa. Upon NaOCl incubation, the protein was observed as a covalent dimer with a molecular mass of 40 kDa. The recombinant protein SydRC114S displayed a similar pattern, with dimer formation under oxidizing conditions. In contrast, the SydRC16S and SydRC16SC114S derivatives remained mostly in the monomeric form. Altogether, these data strongly suggest that NaOCl directly oxidizes SydR and induces the formation of an intermolecular disulfide bond between C16 residues (C16-C16), thus precluding DNA-binding. The role of cysteine residues in DNA-binding was also analyzed (Fig. 3C). EMSA assays showed that SydRC114S was still able to bind to the target DNA, whereas the mutation of the C16 in SydRC16S and SydRC16SC114S impaired DNA-binding. These results indicate that C16 is essential for SydR binding activity.

To strengthen the DNA-binding analysis, a comparison of SMa2023 expression in *S. meliloti* strains overexpressing SydR or SydR derivatives was performed (Fig. 3D). The SMa2023 expression levels in Δ*sydR-*p*trp:sydRC114S* and Δ*sydR*-p*trp*:*sydR* strains were similar and dependent on NaOCl addition. In contrast, gene expression in Δ*sydR-*p*trp:sydRC16S* and Δ*sydR-*p*trp:sydRC16SC114S* was respectively 35-fold and 1000-fold higher compared to untreated Δ*sydR*-p*trp:sydR*, regardless of the redox condition. These data confirm the importance of C16, together with the minor role of C114, in SydR activity.

Overall results show that SydR activity is reversibly inhibited by oxidation, and this redox regulation is mediated by the conserved C16. Moreover, SydR displays a DNA-binding activity that is specifically modulated by NaOCl rather than H_2_O_2_ or tBOOH.

### Analysis of SydR involvement in ROS scavenging in free-living bacteria

As a redox-sensing regulator, SydR could be involved in the modulation of ROS detoxifying pathways. Thus, the effect of SydR inactivation on the sensitivity of *S. meliloti* to NaOCl, H_2_O_2_ and tBOOH was analyzed (Fig. 4). Exogenous concentrations of NaOCl up to 0.2 mM had no effect on growth of the WT strain, while the addition of 0.4 mM NaOCl immediately stopped it. The behavior of Δ*sydR* and Δ*sydR*-p*trp:sydR* strains was not significantly different from that of the WT (Fig. 4A). Moreover, the ability of the three strains to survive after 1h exposure to 10 mM H_2_O_2_ or tBOOH was similar (Fig. 4B). Thus, SydR was not involved in *S. meliloti* resistance to oxidative stress under our conditions.

**FIG 4.**
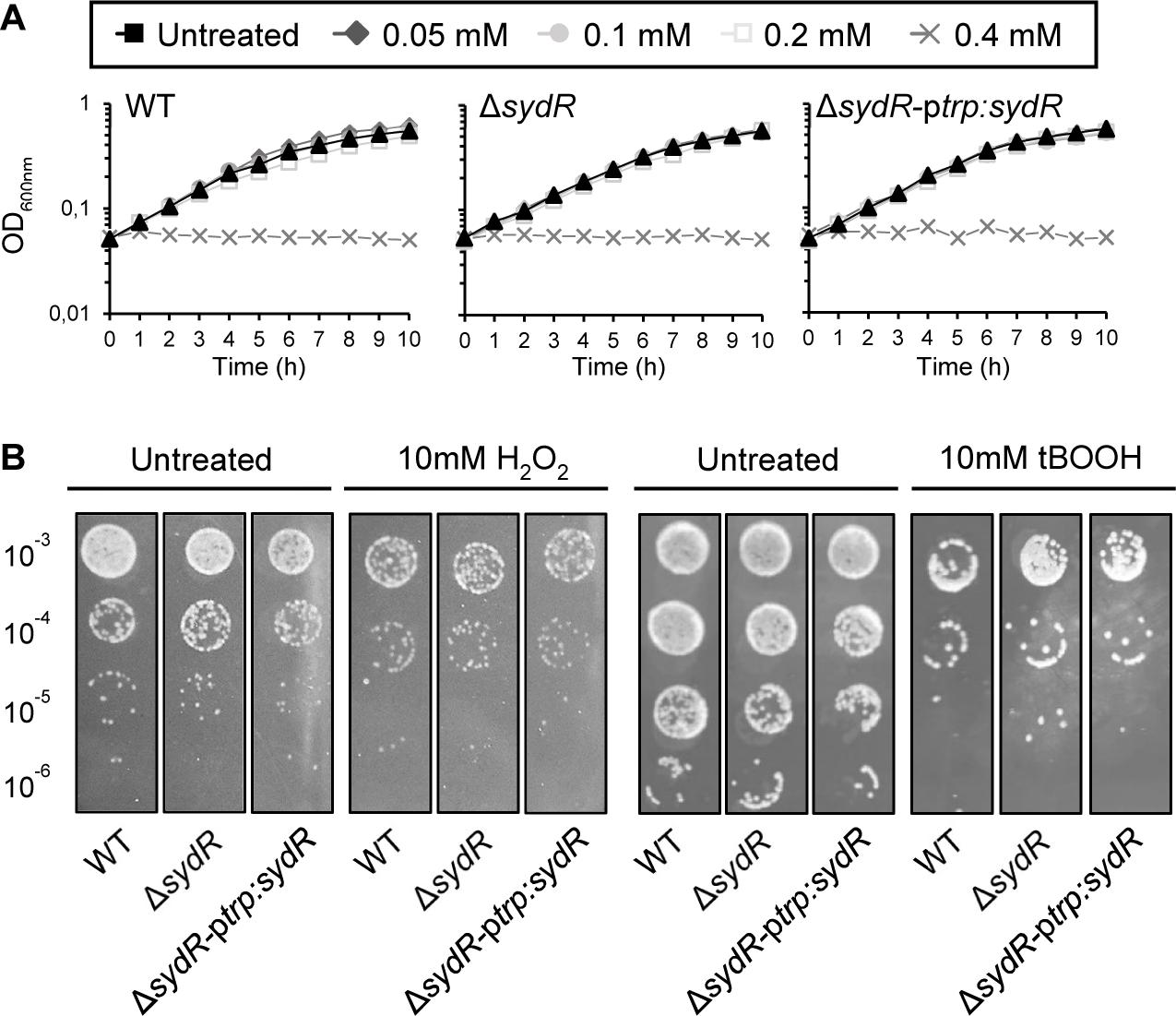
SydR inactivation has no effect on growth or oxidant sensitivity. (A) Growth of WT, Δ*sydR* and Δ*sydR*-p*trp:sydR* strains in M9+CSA in the presence of various concentrations of NaOCl. (B) Bacterial survival after challenge with H_2_O_2_ and tBOOH. WT and Δ*sydR* strains were grown in M9+CSA to OD_600_ of 0.1, then incubated in the presence of either H_2_O_2_ or tBOOH for 1 hour before being serially diluted and spotted on LBMC agar plates. Data are representative of at least three independent experiments.

The effect of *sydR* inactivation on bacteria challenged with ROS was further analyzed by using the genetically encoded biosensor roGFP2-Orp1. This redox probe proved to be an accurate tool for measuring dynamic changes in intracellular H_2_O_2_ pool in *S. meliloti* (8). The roGFP2-Orp1 is directly oxidized by H_2_O_2_, and it can also react with other peroxides and NaOCl, potentially responding to these oxidants in bacterial cultures (22, 23).

The basal oxidation level of roGFP2-Orp1 was similar in WT and ΔsydR strains, corresponding to a highly reducing redox potential EroGFP2 (−290 mV) as previously reported (8). Upon addition of oxidants, similar kinetics of roGFP2-Orp1 oxidation were observed in WT and Δ*sydR* mutant strains. Treatment with H_2_O_2_ (0.1 - 10 mM) resulted in a dose-dependent oxidation of roGFP2-Orp1, followed by biosensor partial recovery within 10 min exposure, as described earlier (Fig. 5A; 8). In cells treated with NaOCl (0.05 - 1 mM) the biosensor also experienced a rapid and reversible oxidation (Fig. 5B). In contrast, roGFP2-Orp1 became highly oxidized within 10 min of adding tBOOH (0.05 - 5 mM), and could recover its reduced state when the treatment time was extended to 1 hour (Fig. 5C; Fig. S2). These results showed the capacity of roGFP2-Orp1 to detect real-time changes in *S. meliloti* intracellular H_2_O_2_, tBOOH or NaOCl levels when cells are exposed to oxidants. However, the dynamics of these pools, together with the toxicity of the molecules were similar in WT and Δ*sydR* strains, revealing no major role of SydR in the control of ROS level in free-living bacteria.

**FIG 5.**
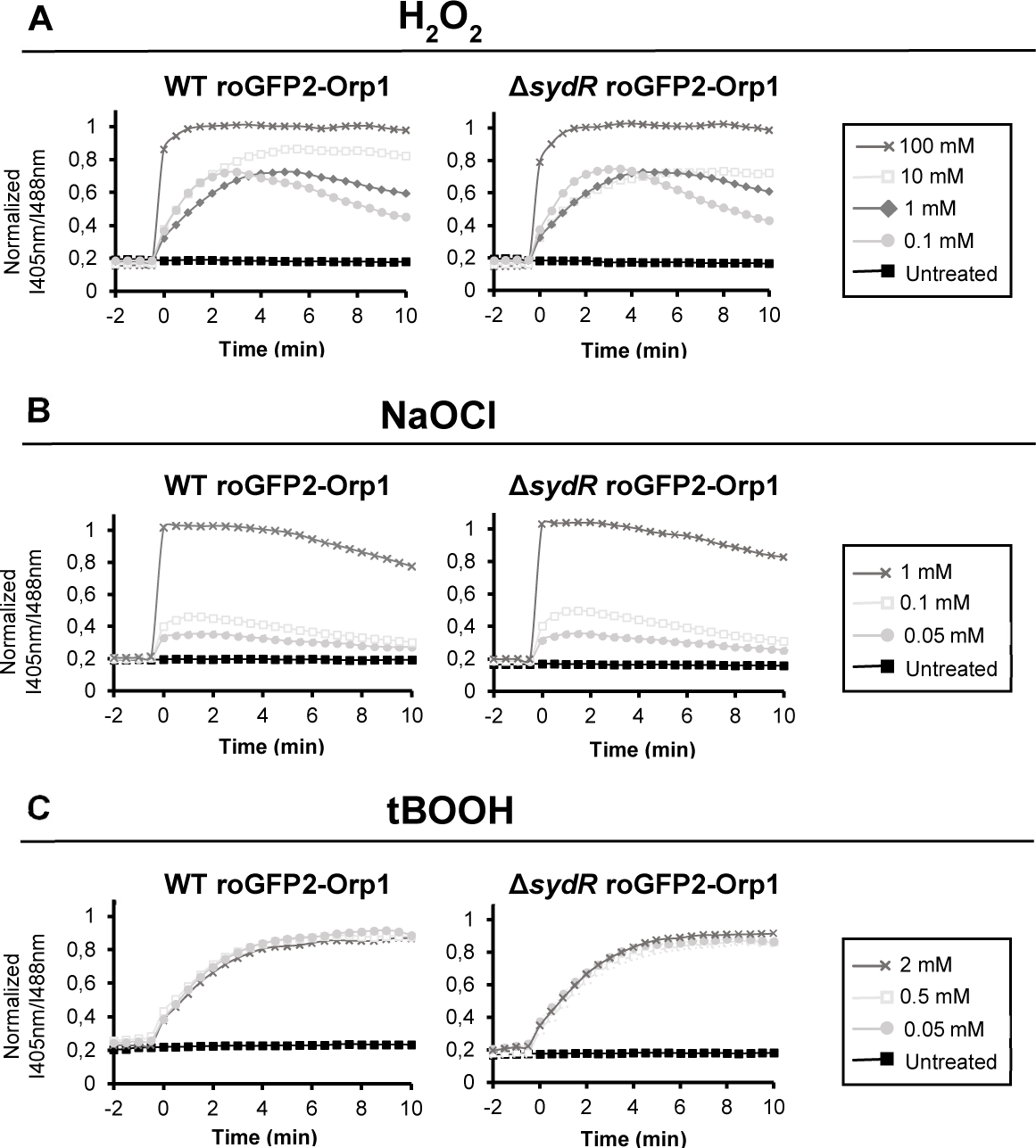
Comparative analysis of WT and Δ*sydR* redox states. Measurements of variations in roGFP2-Orp1 oxidation state in WT and Δ*sydR* treated with various concentrations of H_2_O_2_ (A), tBOOH (B) and NaOCl (C). The biosensor redox state is given by the ratio between fluorescence intensity emission at 405 and 488 nm excitation wavelengths, with a fixed emission of 415 nm (I405 nm/I488 nm), where an increase or decrease of the ration indicates oxidation or reduction, respectively. The maximal and minimal I405 nm/I488 nm ratios, corresponding to fully oxidized and reduced controls were determined in each experiment by cell treatment with 100 mM H_2_O_2_ and 100 mM (DTT) respectively. These values were used to calculate the normalized I405 nm/I488 nm ratios. The values are the means of three independent experiments, with coefficient of variation less than 10%.

### SydR plays a crucial role in the development of *S. meliloti* / *M. truncatula* symbiosis

The symbiotic phenotype of Δ*sydR* mutant was analyzed during interaction with *M. truncatula* (Fig. 6) and *M. sativa* (Fig. S3). Nodulation tests and N_2_ fixation measurements (ARA) were performed using plants inoculated with WT, Δ*sydR* and Δ*sydR-*p*trp:sydR* strains. Inoculation of *M. truncatula* roots with Δ*sydR* mutant led to a drastic reduction of nodule number compared to the WT strain (Nod^−^ phenotype; Fig. 6A). Moreover, most of *M. truncatula* nodules elicited at 21 dpi by the mutant remained white and spherical as non-fixing nodules, while the majority of nodules infected by the WT strain were pink and elongated (Fig. 6B). Consistently, nitrogen fixation in the nodules of roots inoculated with the Δ*sydR* mutant strain was reduced by 75% compared to those infected by the WT strain (Fix^−^ phenotype; Fig. 6C) and the remaining fixation most probably comes from the few pink nodules that have managed to develop (Fig. 6D). The constitutive expression of *sydR* in the Δ*sydR-*p*trp:sydR* strain restored the Nod^+^/Fix^+^ phenotype (Fig. 6C). These data showed that the *sydR* mutation drastically affects nodule development in the symbiotic interaction with *M. truncatula.* To focus on the involvement of SydR in nodule functioning, we constructed a bacterial strain affected in *sydR* expression once released from ITs. This strain (Δ*sydR-*p*nodA:sydR*) carries *sydR* under the control of the p*nodA* promoter that is active from the beginning of interaction to bacterial release inside the plant cell (20). This strain formed nodules where the expression of *sydR* and SMa2023 was respectively down and up regulated compared with WT-induced nodules, as expected (Fig. S3). Moreover, at 21 dpi, the nodules infected by the Δ*sydR-*p*nodA:sydR* strain displayed a Fix^+/−^ phenotype with a 40% deficiency in the acetylene reduction assay (ARA) (Fig. 6E). These results demonstrate that SydR activity contributes to both nodule formation and functioning. In symbiosis with *M. sativa*, roots inoculated by Δ*sydR* also formed less nodules than roots inoculated with WT, but the deficit is less drastic compared to *M. truncatula* roots (reduction of nodule number by 35% *versus* 63%, as compared to WT, at 21 dpi; Fig. S4A). Moreover, the nitrogen fixation capabilities of WT and Δ*sydR* infected nodules were similar (Fig. S4B).

**FIG 6.**
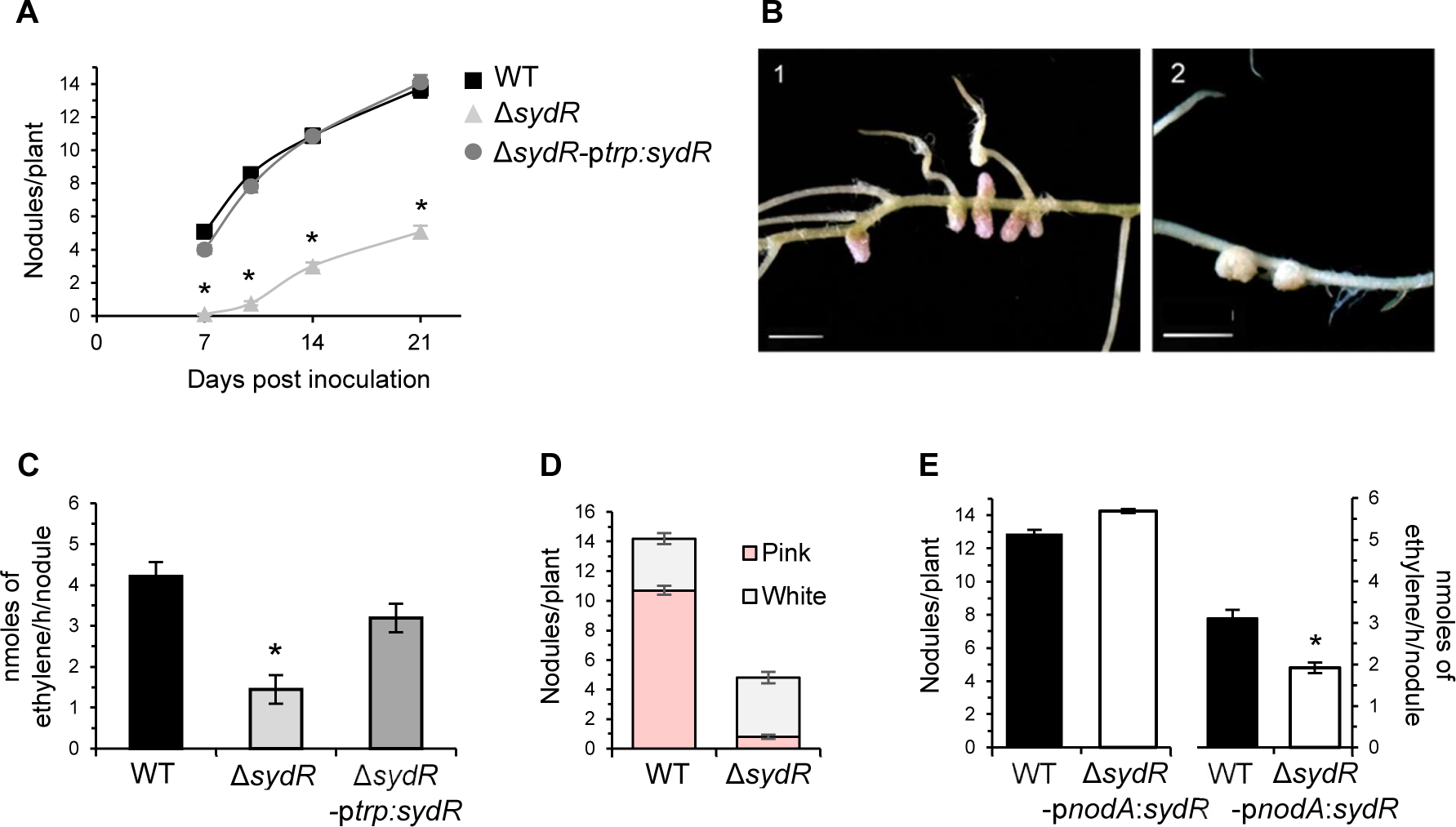
SydR plays a crucial role in *S. meliloti / M. truncatula* symbiosis. (A) Nodulation kinetics of *M. truncatula* plants inoculated with WT, Δ*sydR* and Δ*sydR-*p*trp:sydR* strains. (B) Representative images of *M. truncatula* root nodules after inoculation with WT (1) and Δ*sydR* (2). Scales: 1 cm (1); 0.25 cm (2) (C) Nitrogen fixation activity, determined by acetylene reduction assay (ARA) at 21 dpi. (D) Number of white and pink nodules at 21 dpi. (E) Nodule number and ARA from WT- and Δ*sydR*-p*nodA:sydR*-inoculated roots at 21 dpi. The values shown are the means ± SEM of three independent experiments. Non-parametric Kruskal-Wallis and *post-hoc* Conover-Iman tests with Benjamini-Hochberg correction (A, C), and Mann-Whitney test (E) were used to assess the statistical significance of differences compared to the WT strain (*, p<0.05).

### An essential role of SydR in the infection process

The step of root invasion affected by the *sydR* mutation in *M. truncatula* symbiosis was thereafter specified. Bacterial progression in roots was examined by using strains expressing *lacZ* under the control of a promoter highly expressed *in planta* (p*hemA:lacZ* transcriptional fusion). Roots were inoculated with WT, Δ*sydR* and, for comparison, with an *exoA* mutant known to form early aborted IT (24). In roots inoculated with WT, bacteria were detected within IT at 4 dpi, and readily visible inside nodules at 10 dpi (Fig. 7A.1-2). In the case of roots inoculated with the Δ*sydR* mutant, *lacZ* staining at 4 dpi showed that bacteria mostly accumulate as micro-colonies inside root hair curling (RHC) (Fig. 7A.3). These micro-colonies persisted at 10 dpi, on the surface of small nodule-like bumps devoid of bacteria (Fig. 7A.4). In rare cases, infected nodules were also observed (Fig. 7B). The *exoA* mutant elicited only empty bumps with persistent micro-colonies on the surface (Fig. 7A.5-6). These observations showed that the Δ*sydR* mutant is blocked at the initiation of infection, similar to the *exoA* mutant, once the infection pocket is formed.

**FIG 7.**
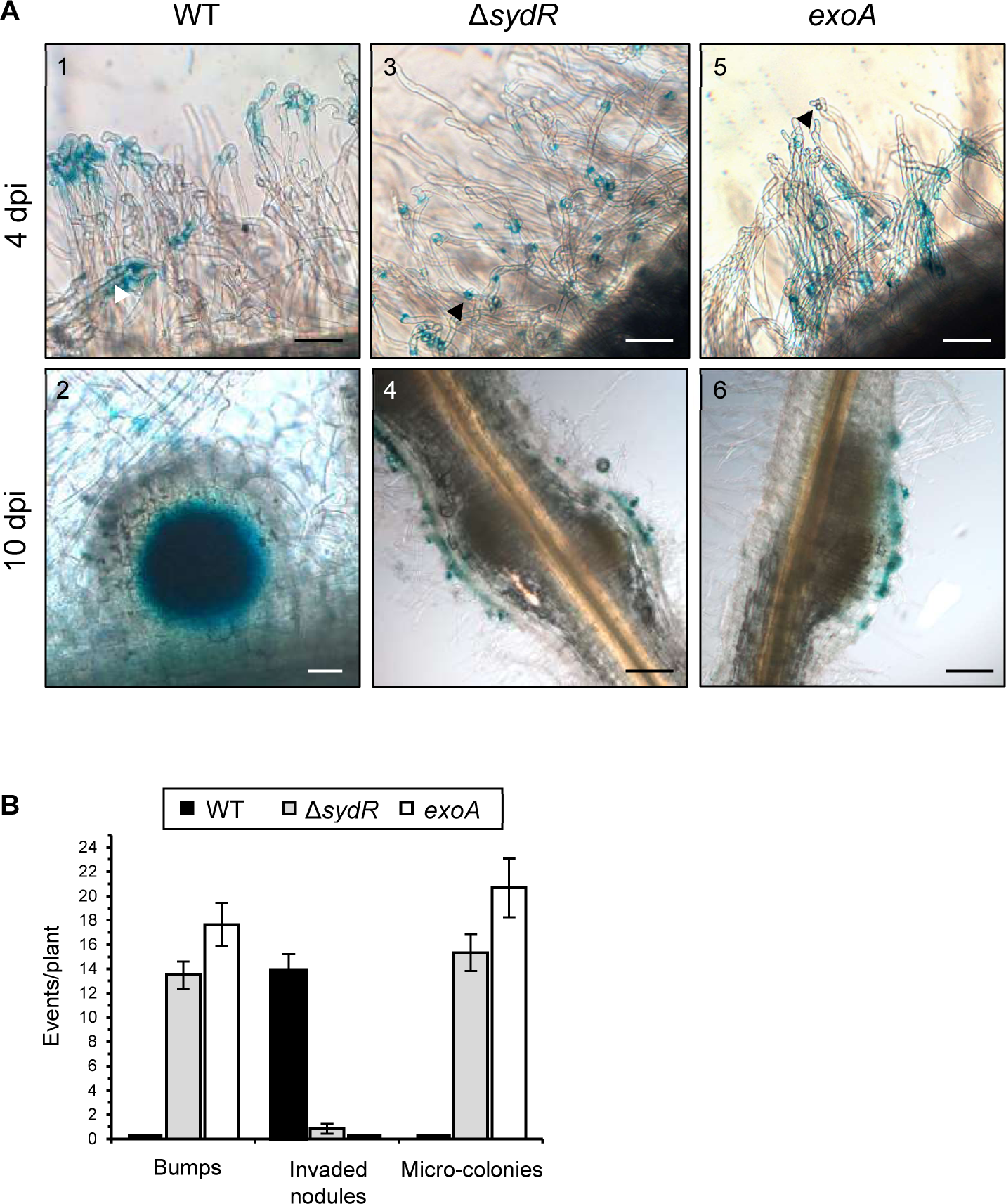
Δ*sydR* mutant is defective in root infection. (A) Light microscopic images of *M. truncatula* roots inoculated with the WT (1, 2), Δ*sydR* (3, 4) and *exoA* (5, 6) strains expressing p*hemA:lacZ* reporter fusion. β-galactosidase activity (blue staining) was detected in *i)* WT bacteria entrapped in root hairs (1), inside infection thread (white arrowhead) and nodule cells (2), and *ii)* Δ*sydR and exoA* bacteria as micro-colonies mainly accumulated in RHCs (black arrowhead; 3, 5) and at the surface of bumps (4, 6). Scales: 50 µm (1, 2, 3, 5); 200 µm (4, 6). (B) Number of bumps, invaded nodules and micro-colonies at 10 dpi. The values shown are the mean ± SEM of three independent experiments.

The response to NFs and initiation of root infection were analyzed using plants expressing the reporter fusion p*MtENOD11-gusA*, and in RT-qPCR experiments. *MtENOD11* is an early nodulin gene expressed firstly in the root epidermis, later in the cortex around the infection threads and then in the central tissue of young nodules (25). As previously described, blue staining corresponding to GUS activity was detected in transgenic roots inoculated with the WT strain *i)* in epidermal cells within 1 dpi, *ii)* restricted to infected root hairs and activated cortical cells within 4 dpi, and *iii)* later in the core tissue of young nodules (Fig. 8A.1-3). By comparison, the expression of *GUS* fusion in roots inoculated with the Δ*sydR* mutant was detected in epidermal cells at 4 dpi, and then in cortical infection zones at 7 dpi (Fig. 8A.4-6). These observations show a delayed induction of *MtENOD11* expression in roots inoculated with the mutant strain, indicating an alteration of the plant response. In parallel, changes in NF-dependent gene expression associated with pre-infection step (*MtNIN*, a key regulator of the NF pathway, in addition to *MtENOD11*), or linked directly to the infection process (*MtLYK3* and *MtN20*) were investigated by RT-qPCR (26–28). As shown in Fig. 8B, *MtNIN* and *MtENOD11* increased upon inoculation with WT and Δ*sydR* mutant strains. However, the induction of both genes occurs significantly later in Δ*sydR* mutant as compared to WT (4 dpi versus 1 dpi). In addition, the expression level of *MtLYK3* decreased, while *MtN20* increased, in roots inoculated with WT strain but remained constant over 7 dpi in those inoculated with the mutant strain.

**FIG 8.**
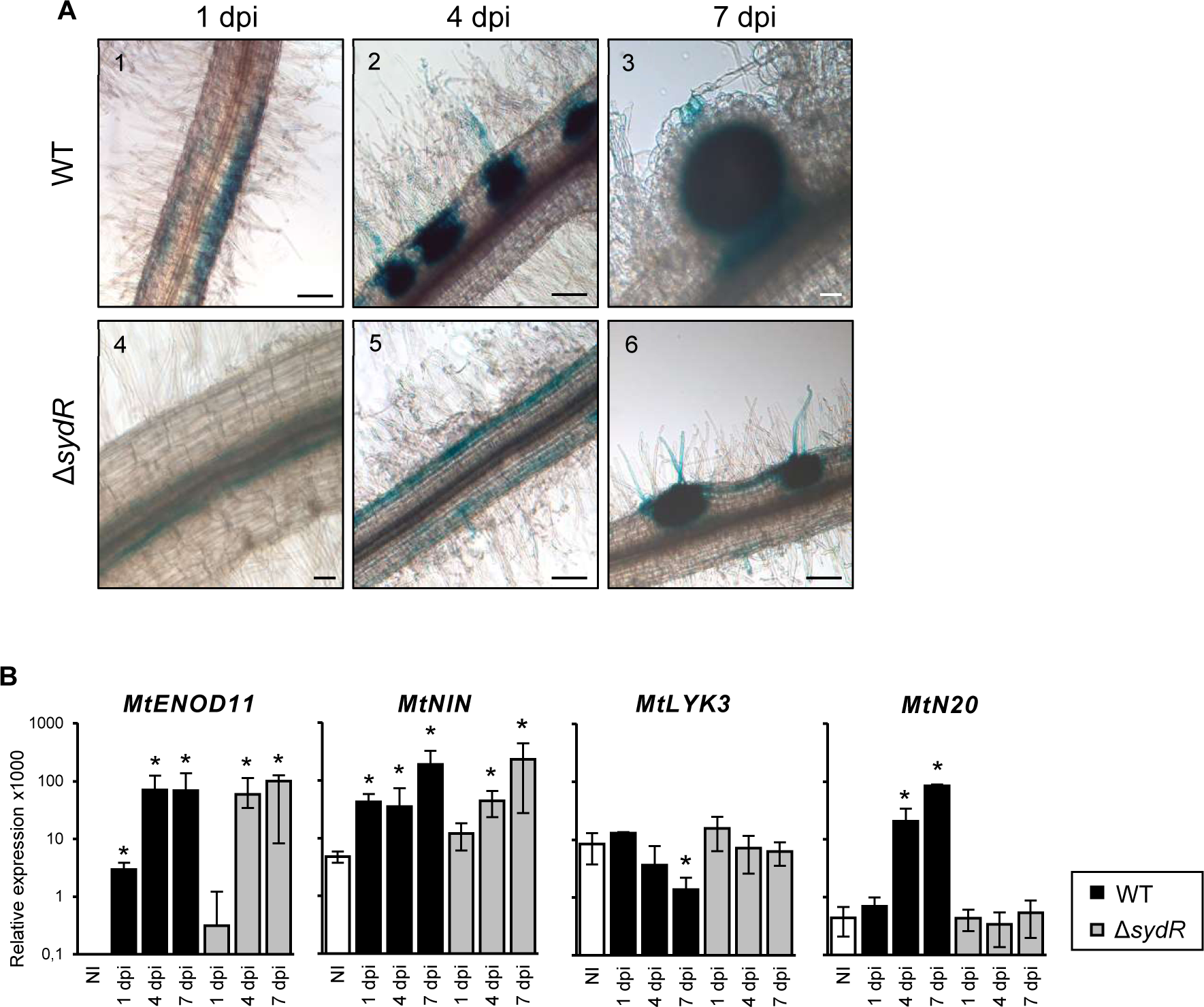
Root inoculation with Δ*sydR* mutant leads to delayed pre-infection and abortive infection events. (A) Light microscopic images of *M. truncatula* transgenic roots expressing p*MtENOD11-gusA* fusion inoculated with the WT and Δ*sydR* strains. GUS activity (blue staining) was visible in epidermal cells in the region of developing root hairs (1, 5), in activated cortical cells corresponding to the initial infection site (2, 6), and in invaded tissues of WT-induced nodules (3). Scales: 50 µm (3, 4); 200 µm (1, 2, 5, 6). (B) RT-qPCR analysis of the expression of *M. truncatula* genes in roots inoculated with either WT or Δ*sydR* at 1, 4 and 7 dpi. The expression ratios for marker genes relative to reference genes are shown. Expression level of genes in non-inoculated roots (NI) was also determined. The values shown are the means ± SEM of three independent experiments. Non-parametric Kruskal-Wallis and *post-hoc* Conover-Iman tests with Benjamini-Hochberg correction were used to assess the statistical significance of differences between inoculated and non-inoculated roots (*, p<0.05).

All together, these data show that the NF-induced signaling pathway is active during the interaction between *M. truncatula* and the Δ*sydR* mutant strain. However, the transcriptional response of the plant and root colonization are altered by the mutation.

## DISCUSSION

In this work, we characterized a new thiol-based redox sensor of *S. meliloti*, SydR, which is critical for the establishment of symbiosis with *M. truncatula*. SydR is one of the 17 proteins annotated as MarR family transcriptional regulators in the *S. meliloti* Genome Database. A high number of MarR-type regulators among others, is a typical feature of bacteria that adopt different lifestyles and have to respond to environmental changes (29, 30). Indeed, several MarR family members have been shown to play a major role in various symbiotic or pathogenic interactions with plants, such as *S. meliloti* WggR and *Dickeya dadantii* PecS (31–34). To a lesser degree, OhrR of *Azorhizobium caulinodans* has also been involved in a symbiotic interaction with host plant (35). SydR is the first redox-sensing MarR-type regulator of that plays a crucial role in the interaction with *M. truncatula*.

Like most MarR-type regulators, SydR represses transcription of the adjacent gene (SMa2023). This repression (∼1000-fold) is variously relieved by addition of NaOCl, H_2_O_2_ and tBOOH. Indeed, in the WT strain, the transcription of SMa2023 was increased ∼100-fold and ∼2-fold upon treatment with NaOCl and H_2_O_2_ / tBOOH respectively, showing that NaOCl is a more potent inducer than hydroperoxides for expression of SydR target gene. In comparison, the expression of OhrR target genes, *ohrA* in *B. subtilis,* and *ohr* in *S. meliloti*, is similarly induced by NaOCl and organic hydroperoxides (this study Fig. S1; 36, 37).

Our data furthermore demonstrated that NaOCl has a direct effect on SydR *in vitro* and leads to its release from the operator DNA. They point out that, between the 2 cysteines of SydR, the conserved C16 alone is essential for redox-sensing and engaged in intermolecular disulfide bond and covalent dimerization of the protein. Up to now, NaOCl was shown to trigger formation of an intermolecular disulfide bond between two distinct cysteines, *i.e.* in *B. subtilis* YodB and HypR, or S-thiolation of the single redox-sensitive cystein of *B. subtilis* OhrR (37–39). The redox-sensing mechanism described here for SydR was reported in a few cases, such as the 1-Cys type QorR of *Corynebacterium glutamicum* and HypS of *Mycobacterium smegmatis* (40,41). This raises the question of whether the two *S. meliloti* SydR paralogs, OhrR and SMa01945 that contain both conserved cysteines, share the same mechanism. These data highlight the role of SydR as a redox-switch under hypochlorite/NaOCl and, potentially, hydroperoxide-induced oxidative stress. The physiological signal regulating SydR activity during interaction remains, however, to be determined. Especially hypochlorite, which is produced by the animal immune system (42), has not been detected in plants.

Thiol-disulfide switches in redox-sensing regulators mainly result in the activation of specific detoxification pathways to restore cellular redox homeostasis upon stress conditions (1). Our data on SydR, both toxicity assays and kinetic analyses of roGFP2-Orp1 oxidation, suggested that the regulator does not play a significant role in the response to exogenous H_2_O_2_, tBOOH or NaOCl. Maybe a difference in ROS sensitivity could be observed by inactivating SydR target(s) rather than *sydR* itself. This has been exemplified by the work of Fontenelle *et al*, (16), where inactivation of *ohr*, but not *ohrR*, modified the sensitivity of *S. meliloti* to oxidant.

SydR is required for the infection of *M. truncatula*. Firstly, the Δ*sydR* mutant gives a drastically reduced number of nodules as compared to WT and generates non-invaded bumps instead. Secondly, microscopic and RT-qPCR analyses showed that early NF signaling, cortical cells activation, and nodule primordium initiation still arise in roots inoculated with Δ*sydR.* Similar observations were obtained with the *exoA* mutant blocked at the stage of IT initiation (our data, 43). Likewise, the repression of *MtLYK3* expression that is directly linked to infection, did not occur in roots inoculated by one or the other mutant (26, 28). Finally, it can be assumed that the infection process in Δ*sydR-*inoculated roots arrests earlier than the process in *exoA-*inoculated roots since, unlike the latter, it lost the ability to trigger *MtN20* induction. Inoculation with the Δ*sydR* mutant, moreover, led to a delayed induction of *MtENOD11* and *MtNIN* expression. Due to lack of infection, most of the cortical cells activated by Δ*sydR* inoculation might be no longer susceptible to signals that induce further nodule development. Finally, by complementing only the infection step, it was possible to show the involvement of SydR in the optimal nodule functioning.

By contrast, Δ*sydR* mutant induces the formation of nitrogen-fixing nodules with *M. sativa*. These data are similar to the results of Zhang *et al*, (17) with a SMa2020 deletion mutant generated in Rm1021 background. They show the importance of plant species (*M. truncatula versus M. sativa)* in the outcome of interaction. Indeed, distinct phenotypic effects across different genetic backgrounds have been already observed such as the formation of non-invaded bumps and effective nodules on, respectively, *M. truncatula* and *M. sativa* roots, inoculated with the same *S. meliloti* lipopolysaccharide mutant (44, 45). As another example, inactivation of the *relA* gene involved in the stringent response, was shown to affect the capacity of *S. meliloti* to infect *M. sativa* only (46). Thus, it can be assumed that checkpoints involved in bacterial root infection function differentially in *M. truncatula* and *M. sativa*. Most likely, genotypic and phenotypic analysis of bacteria isolated from the rare pink nodules produced by Δ*sydR* in *M. truncatula* roots will help identifying functions specifically involved in *S. meliloti / M. truncatula* interaction.

Further studies will be required to identify the mechanisms/targets controlled by SydR, particularly those whose fine regulation is required for the symbiosis establishment and functioning.

## MATERIALS AND METHODS

### Bacterial strains and growth conditions

All bacterial strains and plasmids used in this study are listed in Table S2. *Escherichia coli* strains were grown at 37°C in Luria-Bertani (LB) medium. *S. meliloti* strains were grown at 30°C in LB medium supplemented with 2.5 mM MgSO_4_ and 2.5 mM CaCl_2_ (LBMC), in M9 medium and in M9-CSA with appropriate antibiotics, as specified in text S1 in the Supplemental Material.

### Preparation of bacterial samples for RT-qPCR analysis and toxicity tests

Cultures of *S. meliloti* strains were grown in M9 to mid-log phase (OD_600_ ∼0.3), then divided into 10 ml aliquots and exposed to various oxidants for 10 min (1 mM H_2_O_2_; 200 μM tBOOH; 50 μM Plumbagin; 0.2 mM NaOCl) or 30 min (25 μM Spermine NONOate). The bacterial cultures were immediately centrifuged, and pellets were frozen at *−*80°C until RNA extraction. For testing oxidant toxicity, cultures grown to mid-log phase in M9-CSA were diluted to an OD_600_ of ∼ 0.1, then divided into 5 ml aliquots that were incubated with 1 mM H_2_O_2_ or 10 mM tBOOH for 1 h, or further grown with various concentrations of NaOCl. Samples treated with H_2_O_2_ or tBOOH were serially diluted, and 10 µl were spotted in three replicates on LBMC plates; CFU were counted after 48 h of incubation at 30°C. The effect of NaOCl was evaluated by monitoring OD_600_ for 6h. An untreated culture was included as a control in each experiment.

### Plant growth conditions

*M. truncatula* ecotype *Jemalong* A17 and *M. sativa* L. var. Europe (alfalfa) were used as the host plants to test nodulation and nitrogen fixation of *S. meliloti* strains. Seed germination and plant growth procedures were performed as described previously (47, 48), with modifications summarized in text S1.

### Construction of *S. meliloti* mutants

The ΔSMc03824, SMc00146 and Δ*sydR* mutants were constructed using allelic exchange mutagenesis, as described in text S1.

### Molecular cloning and mutagenesis of *sydR*

Primers used for DNA amplification were listed in Table S3. Molecular cloning and mutagenesis of *sydR* were performed as described in text S1.

### Real-time measurements of intracellular redox potential

Variations in intracellular redox potential of *S. meliloti* were monitored with ratiometric roGFP2 fluorescence measurements (excitation at 405 and 488 nm wavelenghts, emission at 515 nm) using a spectrofluorimeter luminometer (Xenius, Safas, Monaco), as previously described (8).

### RNA extraction and RT*-*qPCR assays

RNA extraction and analysis were performed as described in text S1.

### Purification of SydR wild-type and mutant proteins

The production and purification of recombinant proteins were performed as described in text S1.

### Intersubunit disulphide bond assay

SydR’ and mutant proteins were reduced in 100mM DTT for 2h and desalted using ultra centrifugal columns with 10 NMWL. The proteins were oxidized by treatment with 100mM NaCl for 30 min. Additionaly, 100 mM N-ethylmaleimide (NEM) was applied to each sample for 2h at room temperature to reduce the formation of non-specific disulfide bonds. Finally, the samples were analyzed by 10% non-reducing-SDS-page.

### Electrophoretic mobility shift assay (EMSA)

For EMSA analysis, the 140 bp DNA fragment covering the *sydR*-SMa2023 intergenic region was amplified using *sma2020*_23F/*sma2020*_23R primers. DNA binding reactions were implemented in binding buffer (10 mM Tris-HCl (pH8), 50 mM KCl, 0,5 mM EDTA, 10% glycerol) containing various amounts of SydR’ WT protein and mutant variants, and 10 nM DNA. The samples were incubated at 25°C for 25 min and loaded on 8% polyacrylamide gel in 0,5X Tris borate-EDTA buffer. The Diamond™ Nucleic Acid Dye (Promega) staining was used to visualize DNA on the gel. The gel was analyzed under UV light (UVIDOC HD6, Cambridge).

### Nitrogen fixation assays

N_2_-fixation activity was determined at 21 dpi by assessing C_2_H_2_ reduction using gas chromatography (Agilent Technologies 6890N) as previously described (48). At least 50 plants from three independent biological repetitions were analyzed for each inoculation.

### Microscopy and histology analysis

Roots expressing the p*MtENOD11-gusA* fusion were harvested at 1, 4, and 7 dpi (n = 4 roots per time point). Roots inoculated with bacterial strains expressing the p*hemA:lacZ* fusion were harvested at 4 and 10 dpi (n=6 roots per time point), or 14 dpi. GUS and β-galactosidase staining as described previously (25, 49), with modifications summarized in text S1. Stained roots and nodule sections were observed under a transmission light microscope (Zeiss Axioplan II).

## ACKNOWLEDGMENTS

We are grateful to Marc Bosseno for assistance with plant experiments. We also thank Claude Castella, Jean-Luc Gatti and Nicolas Pauly for helpful discussions. The present work has benefited from the Cell Imagery Platform (ISC PlantBIOs) facility of the Institute Sophia Agrobiotech (ISA), with Olivier Pierre assistance.

FN was supported by a Ph.D. scholarship from the Ministère de l’Enseignement Supérieur, de la Recherche et de l’Innovation (MESRI). DF was supported by a sabbatical grant from Azarbaijan Shahid Madani University. The laboratory was the recipient of a collaborative funding between INRAE and the Azarbaijan Shahid Madani University, as part of the Gundishapur program between the French Ministère de l’Europe et des Affaires étrangères and MESRI, and the Iranian Ministry of Science, Research and Technology (PHC GUNDISHAPUR - 42938YA). This work received financial support from Institut National de Recherche pour l’Agriculture, l’Alimentation et l’Environnement (INRAE), the Centre National de la Recherche Scientifique (CNRS), Université Côte d’Azur (UCA), the French Government (National Research Agency, ANR) through the “Investments for the Future” of the Labex Signalife (ANR-11-LABX-647-0028) and the IDEX UCA JEDI (ANR-15-IDEX-01).

PF, GA, and KM conceived and designed research; FN, DF, JM, MP, AC, and GA performed research; FN, DF, JM, AC, GA, and KM analyzed data; and FN, DF, PF, GA, and KM wrote the paper.

## REFERENCES

1. Antelmann H, Helmann JD. 2011. Thiol-based redox switches and gene regulation. Antioxid. Redox Signal. 14:1049–1063. 10.1089/ars.2010.3400.

2. Damiani I, Pauly N, Puppo A, Brouquisse R, Boscari A. 2016. Reactive oxygen species and nitric oxide control early steps of the legume – *Rhizobium* symbiotic interaction. Front Plant Sci 7:1–8. 10.3389/fpls.2016.00454.

3. Syska C, Brouquisse R, Alloing G, Pauly N, Frendo P, Bosseno M, Dupont L, Boscari A. 2019. Molecular weapons contribute to intracellular rhizobia accommodation within legume host cell. Front Plant Sci. Frontiers Media S.A. 10.3389/fpls.2019.01496.

4. Camejo D, Guzmán-Cedeño A, Vera-Macias L, Jiménez A. 2019. Oxidative post-translational modifications controlling plant-pathogen interaction. Plant Physiology and Biochemistry. Elsevier Masson SAS 10.1016/j.plaphy.2019.09.020.

5. Soumare A, Diedhiou AG, Thuita M, Hafidi M, Ouhdouch Y, Gopalakrishnan S, Kouisni L. 2020. Exploiting biological nitrogen fixation: A route towards a sustainable agriculture. Plants. MDPI AG 10.3390/plants9081011.

6. Oldroyd GED, Murray JD, Poole PS, Downie JA. 2011. The rules of engagement in the legume-rhizobial symbiosis 10.1146/annurev-genet-110410-132549.

7. Ribeiro CW, Alloing G, Mandon K, Frendo P. 2015. Redox regulation of differentiation in symbiotic nitrogen fixation. Biochim Biophys Acta Gen Subj. Elsevier B.V. 10.1016/j.bbagen.2014.11.018.

8. Pacoud M, Mandon K, Cazareth J, Pierre O, Frendo P, Alloing G. 2022. Redox-sensitive fluorescent biosensors detect *Sinorhizobium meliloti* intracellular redox changes under free-living and symbiotic lifestyles. Free Radic Biol Med 184:185–195. 10.1016/j.freeradbiomed.2022.03.030.

9. Mandon K, Nazaret F, Farajzadeh D, Alloing G, Frendo P. 2021. Redox Regulation in diazotrophic bacteria in interaction with plants. Antioxidants 10:880. 10.3390/antiox10060880.

10. Hillion M, Antelmann H. 2015. Thiol-based redox switches in prokaryotes. Biol Chem. 396:415–4441. 10.1515/hsz-2015-0102

11. Jamet A, Kiss E, Batut J, Puppo A, Hérouart D. 2005. The *katA* catalase gene is regulated by OxyR in both free-living and symbiotic *Sinorhizobium meliloti*. J Bacteriol 187:376–381. 10.1128/JB.187.1.376-381.2005.

12. Lehman AP, Long SR. 2018. OxyR-dependent transcription response of *Sinorhizobium meliloti* to oxidative stress. J Bacteriol 200:e00622–17. 10.1128/JB.00622-17.

13. Luo L, Yao SY, Becker A, Rüberg S, Yu GQ, Zhu JB, Cheng HP. 2005. Two new *Sinorhizobium meliloti* LysR-Type transcriptional regulators required for nodulation. J Bacteriol 187:4562–4572. 10.1128/JB.187.13.4562-4572.2005.

14. Tang GR, Lu DW, Wang D, Luo L. 2013. *Sinorhizobium meliloti lsrB* is involved in alfalfa root nodule development and nitrogen-fixing bacteroid differentiation. Chinese Science Bulletin 58:4077–4083. 10.1007/s11434-013-5960-6.

15. Deochand DK, Grove A. 2017. MarR family transcription factors: dynamic variations on a common scaffold. Crit Rev Biochem Mol Biol. Taylor and Francis Ltd 10.1080/10409238.2017.1344612.

16. Fontenelle C, Blanco C, Arrieta M, Dufour V, Trautwetter A. 2011. Resistance to organic hydroperoxides requires *ohr* and *ohrR* genes in *Sinorhizobium meliloti*. BMC Microbiology 11:100. 10.1186/1471-2180-11-100.

17. Zhang L, Li N, Wang Y, Zheng W, Shan D, Yu L, Luo L. 2022. *Sinorhizobium meliloti ohrR* genes affect symbiotic performance with alfalfa (*Medicago sativa*). Environ Microbiol Rep 14:595–603. 10.1186/1471-2180-11-100.

18. Barloy-Hubler F, Chéron A, Hellégouarch A, Galibert F. 2004. Smc01944, a secreted peroxidase induced by oxidative stresses in *Sinorhizobium meliloti* 1021. Microbiology (N Y) 150:657–664. 10.1099/mic.0.26764-0.

19. Bahlawane C, Baumgarth B, Serrania J, Rüberg S, Becker A. 2008. Fine-Tuning of galactoglucan biosynthesis in *Sinorhizobium meliloti* by differential WggR (ExpG)-, PhoB-, and MucR-dependent regulation of two promoters. J Bacteriol 190:3456–3466. 10.1128/JB.00062-08

20. Roux B, Rodde N, Jardinaud MF, Timmers T, Sauviac L, Cottret L, Carrère S, Sallet E, Courcelle E, Moreau S, Debellé F, Capela D, De Carvalho-Niebel F, Gouzy J, Bruand C, Gamas P. 2014. An integrated analysis of plant and bacterial gene expression in symbiotic root nodules using laser-capture microdissection coupled to RNA sequencing. Plant Journal 77:817–837. 10.1111/tpj.12442

21. Wilkinson SP, Grove A. 2006. Ligand-responsive transcriptional regulation by members of the MarR family of winged helix proteins. Curr Issues Mol Biol 8:51–62. 10.21775/cimb.008.051. 10.1111/tpj.12442.

22. Müller A, Schneider JF, Degrossoli A, Lupilova N, Dick TP, Leichert LI. 2017. Systematic *in vitro* assessment of responses of roGFP2-based probes to physiologically relevant oxidant species. Free Radic Biol Med 106:329–338. 10.1016/j.freeradbiomed.2017.02.044

23. Degrossoli A, Müller A, Xie K, Schneider JF, Bader V, Winklhofer KF, Meyer AJ, Leichert LI. 2018. Neutrophil-generated HOCl leads to non-specific thiol oxidation in phagocytized bacteria. Elife 7:e32288. 10.7554/eLife.32288.

24. Mitra RM, Long SR. 2004. Plant and Bacterial Symbiotic Mutants define three transcriptionally distinct stages in the development of the *Medicago truncatula*/*Sinorhizobium meliloti* Symbiosis. Plant Physiol 134:595–604. 10.1104/pp.103.031518.

25. Journet E-P, El-Gachtouli N, Vernoud V, De Billy F, Pichon M, Dedieu A, Arnould C, Morandi D, Barker DG, Gianinazzi-Pearson V. 2001. *Medicago truncatula* ENOD11: A Novel RPRP-Encoding Early Nodulin Gene Expressed During Mycorrhization in Arbuscule-Containing Cells. Mol Plant Microbe Interact. 14:737–748. 10.1094/MPMI.2001.14.6.737

26. Smit P, Limpens E, Geurts R, Fedorova E, Dolgikh E, Gough C, Bisseling T. 2007. *Medicago* LYK3, an entry receptor in rhizobial nodulation factor signaling. Plant Physiol 145:183–191. 10.1104/pp.107.100495.

27. Marsh JF, Rakocevic A, Mitra RM, Brocard L, Sun J, Eschstruth A, Long SR, Schultze M, Ratet P, Oldroyd GED. 2007. *Medicago truncatula* NIN is essential for rhizobial-independent nodule organogenesis induced by autoactive calcium/calmodulin-dependent protein kinase. Plant Physiol 144:324–335. 10.1104/pp.106.093021.

28. Moreau S, Verdenaud M, Ott T, Letort S, De Billy F. 2011. Transcription reprogramming during root nodule development in *Medicago truncatula*. PLoS One 6:16463. 10.1371/journal.pone.0016463.

29. Gupta A, Pande A, Sabrin A, Thapa SS, Gioe BW, Grove A. 2019. MarR family transcription factors from *Burkholderia* Species: Hidden clues to control of virulence-associated genes. Microbiol Mol Biol Rev 83:e00039–18. 10.1128/MMBR.00039-18.

30. Grove A. 2017. Regulation of metabolic pathways by MarR family transcription factors. Comput Struct Biotechnol J 15:366–371. 10.1016/j.csbj.2017.06.001.

31. Hommais F, Oger-Desfeux C, Van Gijsegem F, Castang S, Ligori S, Expert D, Nasser W, Reverchon S. 2008. PecS is a global regulator of the symptomatic phase in the phytopathogenic bacterium *Erwinia chrysanthemi* 3937. J Bacteriol 190:7508–7522. 10.1128/JB.00553-08.

32. McIntosh M, Krol E, Becker A. 2008. Competitive and cooperative effects in quorum-sensing-regulated galactoglucan biosynthesis in *Sinorhizobium meliloti*. J Bacteriol 190:5308–5317. 10.1128/JB.00063-08.

33. Mueller K, González JE. 2011. Complex regulation of symbiotic functions is coordinated by MucR and quorum sensing in *Sinorhizobium meliloti*. J Bacteriol 193:485–496. 10.1128/JB.01129-10.

34. Pédron J, Chapelle E, Alunni B, Van Gijsegem F. 2018. Transcriptome analysis of the *Dickeya dadantii* PecS regulon during the early stages of interaction with *Arabidopsis thaliana*. Mol Plant Pathol 19:647–663. 10.1111/mpp.12549.

35. Si Y, Guo D, Deng S, Lu X, Zhu J, Rao B, Cao Y, Jiang G, Yu D, Zhong Z, Zhu J. 2020. Ohr and OhrR are critical for organic peroxide resistance and symbiosis in *Azorhizobium caulinodans* ORS571. Genes (Basel) 11:335. 10.3390/genes11030335.

36. Fuangthong M, Atichartpongkul S, Mongkolsuk S, Helmann JD. 2001. OhrR is a repressor of *ohrA*, a key organic hydroperoxide resistance determinant in *Bacillus subtilis*. J Bacteriol 183:4134–4141. 10.1128/JB.183.

37. Khanh Chi B, Gronau K, Mä der U, Hessling B, rte Becher D, Antelmann H. 2011. S-bacillithiolation protects against hypochlorite stress in Bacillus subtilis as revealed by transcriptomics and redox Proteomics. Mol Cell Prot 10:1–21. 10.1074/mcp.M111.009506.

38. Chi BK, Kobayashi K, Albrecht D, Hecker M, Antelmann H. 2010. The paralogous MarR/DUF24-family repressors YodB and CatR control expression of the catechol dioxygenase CatE in *Bacillus subtilis*. J Bacteriol 192:4571–4581. 10.1128/JB.00409-10.

39. Palm GJ, Khanh Chi B, Waack P, Gronau K, Becher D, Albrecht D, Hinrichs W, Read RJ, Antelmann H. 2012. Structural insights into the redox-switch mechanism of the MarR/DUF24-type regulator HypR. Nucleic Acids Res 40:4178–4192. 10.1093/nar/gkr1316.

40. Ehira S, Ogino H, Teramoto H, Inui M, Yukawa H. 2009. Regulation of quinone oxidoreductase by redox-sensing transcriptional regulator QorR in *Corynebacterium glutamicum*. J Biol Chem 284:16736–16742. 10.1074/jbc.M109.009027.

41. Ngoc Tung Q, Busche T, Van Loi V, Kalinowski J, Antelmann H. 2019. The redox-sensing MarR-type repressor HypS controls hypochlorite and antimicrobial resistance in *Mycobacterium smegmatis*. Free Rad Biol Med 10.1016/j.freeradbiomed.2019.12.032.

42. Andrés CMC, Pérez de la Lastra JM, Juan CA, Plou FJ, Pérez-Lebeña E. 2022. Hypochlorous acid chemistry in mammalian cells–influence on infection and role in various pathologies. Int J Mol Sci 23:10735. 10.3390/ijms231810735.

43. Battisti L, Jimmie L., Leigh L. 1992. Specific oligosaccharide form of the *Rhizobium meliloti* exopolysaccharide promotes nodule invasion in alfalfa. Proc Natl Acad Sci USA 89:5625–5629. 10.1073/pnas.89.12.5625.

44. Nicoud Q, Barrière Q, Busset N, Dendene S, Travin D, Bourge M, Bars L, Boulogne C, Lecroel M, Jenei S, Kereszt A, Kondorosi É, Biondi EG, Timchenko T, Alunni B, Mergaert P. 2021. *Sinorhizobium meliloti* functions required for resistance to antimicrobial NCR peptides and bacteroid differentiation. mBio 12:1–18. 10.1128/mbio.00895-21.

45. Niehaus K, Lagares A, Pühler A. 1998. A *Sinorhizobium meliloti* lipopolysaccharide mutant induces effective nodules on the host plant *Medicago sativa* (Alfalfa) but fails to establish a symbiosis with *Medicago truncatula*. Mol Plant Microbe Interact 11:906–914. 10.1094/MPMI.1998.11.9.906.

46. Wippel K, Long SR. 2019. Symbiotic performance of *Sinorhizobium meliloti* lacking ppGpp depends on the *Medicago* host species. Mol Plant Microbe Interact 32:717–728. 10.1094/MPMI-11-18-0306-R.

47. Pucciariello C, Innocenti G, van de Velde W, Lambert A, Hopkins J, Clément M, Ponchet M, Pauly N, Goormachtig S, Holsters M, Puppo A, Frendo P. 2009. (Homo)glutathione depletion modulates host gene expression during the symbiotic interaction between *Medicago truncatula* and *Sinorhizobium meliloti*. Plant Physiol 151:1186–1196. 10.1104/pp.109.142034.

48. Boncompagni E, Dupont L, Mignot T, Østeräs M, Lambert A, Poggi M-C, Le Rudulier D. 2000. Characterization of a Sinorhizobium meliloti ATP-Binding Cassette Histidine Transporter Also Involved in Betaine and Proline Uptake. J Bac 182:3717–3725. 10.1128/JB.182.13.3717-3725.2000.

49. Boivin C., Camut S., Malpica C., Truchet G., Rosenberg C. 1990. *Rhizobium meliloti* genes encoding catabolism of trigonelline are induced under symbiotic conditions. Plant Cell 2:1157–1170. 10.1105/tpc.2.12.1157.

50. Waterhouse A, Procter J, Martin D, Clamp M, Barton G. 2009. Jalview version 2−a multiple sequence alignment editor and analysis workbench. Bioinformatics 25:1189–1191. 10.1093/bioinformatics/btp033.

