## supplemental data for "SydR, a redox-sensing MarR-type regulator of *Sinorhizobium meliloti*, is crucial for symbiotic infection of *Medicago truncatula* roots"

**Table S1. *S. meliloti* MarR-type regulators that contain cysteine(s).** * corresponds to the number of total reads from laser-capture microdissection of nodules cells coupled to RNA-seq, given in (1). Accession numbers are from the *S. meliloti* genome database (http://sequence.toulouse.inra.fr/meliloti.html).

| Gene (product) | Gene expression in nodule* | Protein length (in aa) | Cystein(s) | References |
|---|---|---|---|---|
| SMa2020 (SydR) | 1122 | 155 | C16, C114 | This study; 2 |
| SMb21317 (WggR) | 1462 | 194 | C148 | 3 |
| SMc00098 (OhrR) | 4853 | 158 | C23, C132 | 4 |
| SMc00146 | 940 | 173 | C30, C139 | This study |
| SMc00384 | 1720 | 169 | C122 | This study |
| SMc00562 | 157 | 150 | C135 | This study |
| SMc01908 | 5013 | 157 | C48, C55 | This study |
| SMc01945 (Cpo) | 487 | 150 | C8 | 5 |
| SMc03824 | 4529 | 159 | C149 | This study |
| SMc04052 | 112 | 148 | C19, C21 | This study |

### REFERENCES


1. Roux B, Rodde N, Jardinaud MF, Timmers T, Sauviac L, Cottret L, Carrère S, Sallet E, Courcelle E, Moreau S, Debellé F, Capela D, De Carvalho-Niebel F, Gouzy J, Bruand C, Gamas P. 2014. An integrated analysis of plant and bacterial gene expression in symbiotic root nodules using laser-capture microdissection coupled to RNA sequencing. Plant Journal 77:817–837. https://doi.org/10.1111/tpj.12442.

2. Zhang L, Li N, Wang Y, Zheng W, Shan D, Yu L, and Luo L. 2022. *Sinorhizobium meliloti ohrR* genes affect symbiotic performance with alfalfa (*Medicago sativa*). Environ Microbiol Rep 14:595–603. https://doi.org/10.1111/1758-2229.13079.

3. Bahlawane C, Baumgarth B, Serrania J, Rüberg S, Becker A. 2008. Fine-tuning of galactoglucan biosynthesis in *Sinorhizobium meliloti* by differential WggR (ExpG)-, PhoB-, and MucR-dependent regulation of two Promoters. J Bacteriol 190:3456–66. https://doi.org/10.1128/JB.00062-08.

4. Fontenelle C, Blanco C, Arrieta M, Dufour V, Trautwetter A. 2011. Resistance to organic hydroperoxides requires *ohr* and *ohrR* genes in *Sinorhizobium meliloti*. BMC Microbiol 11:100. https://doi.org/10.1186/1471-2180-11-100.

5. Barloy-Hubler F, Chéron A, Hellégouarch A, Galibert F. 2004. Smc01944, a secreted peroxidase induced by oxidative stresses in *Sinorhizobium meliloti* 1021. Microbiol 150:657–64. https://doi.org/10.1099/mic.0.26764-0.

**Table S2. Bacterial strains and plasmids used in this study.**

| Strains | Relevant charateristics | Source or reference |
|---|---|---|
| *Escherichia coli* | | |
| DH5α | F- *supE44 ΔlacU169 (Φ80lacZΔM15) hsdR17 recA1 endA1 gyrA96 thi-1* | Invitrogen |
| BL21 (DE3) pLysS | *ompT gal lon hsdSB(rb-mb-) λ(DE3 [lacI lacUV5-T7p07 ind1 sam7 nin5]) [malB+]K-12(λS) pLysS[T7p20 orip15A)*; Cm$^R$ | Invitrogen |
| MT616 | helper strain used for triparental conjugation*, pro-82 thi-1 hsdR17 supE44 reca56 (pRK600)*; Cm$^R$ | 1 |
| SmPI_S17-1.123.03G6 | S17-1 strain carrying plasmid with an SMc00146 internal fragment, used for insertional inactivation of SMc00146 | A. Becker, Germany. |
| *Sinorhizobium meliloti* | | |
| Rm2011 | Wild-type strain, SU47 derivative; Sm$^R$ | 2 |
| Rm2011-pTB° | Rm2011 carrying the pTB° plasmid; Sm$^R$, Spc$^R$ | 3 |
| Rm2011-pTBroGFP2-Orp1 | Rm2011 carrying the pTBroGFP2-Orp1 plasmid; Sm$^R$, Spc$^R$ | 3 |
| Rm2011-pTBGrx1-roGFP2 | Rm2011 carrying the pTBGrx1-roGFP2 plasmid; Sm$^R$, Spc$^R$ | 3 |
| Rm2011-pXLGD4 | Rm2011 carrying the pXLGD4 plasmid; Sm$^R$, Tc$^R$ | 4 |
| Δ*sydR* | Rm2011 derivative carrying an deletion-insertion of SMa2020 (*sydR)*; Sm$^R$, Tc$^R$ | This study |
| Δ*sydR*-pTB° | Δ*sydR* carrying the pTB° plasmid; Sm$^R$, Spc$^R$, Tc$^R$ | This study |
| Δ*sydR*-pTBroGFP2-Orp1 | Δ*sydR* carrying the pTB roGFP2-Orp1 plasmid; Sm$^R$, Spc$^R$, Tc$^R$ | This study |
| Δ*sydR*-pTBGrx1-roGFP2 | Δ*sydR* carrying the pTB Grx1-roGFP2 plasmid; Sm$^R$, Spc$^R$, Tc$^R$ | This study |
| Δ*sydR*-pXLGD4 | Δ*sydR* carrying the pXLGD4 plasmid; Sm$^R$, Tc$^R$ | This study |
| Δ*sydR*-p*trp:sydR* | Δ*sydR rhaS*::pCAP97-SMa2020(WT); Sm$^R$, Tc$^R$, Spc$^R$ | This study |
| Δ*sydR-sydRC16S* | Δ*sydR rhaS*::pCAP97-SMa2020_C16S; Sm$^R$, Tc$^R$, Spc$^R$ | This study |
| Δ*sydR-sydRC114S* | Δ*sydR rhaS*::pCAP97-SMa2020_C114S; Sm$^R$, Tc$^R$, Spc$^R$ | This study |
| Δ*sydR-sydRC16S-C114S* | Δ*sydR rham*:: pCAP97-SMa2020_C16S&C114S; Sm$^R$, Tc$^R$, Spc$^R$ | This study |
| Δ*sydR*-p*nodA*:*sydR* | Δ*sydR rhaS*::pCAP97-SMa2020*(nodA)*; Sm$^R$, Tc$^R$, Spc$^R$ | This study |
| ΔSMc03824 | Rm2011 derivative carrying a deletion of SMc03824; Sm$^R$ | This study |
| SmPI_1021.12.01A9 | Rm1021 derivative with insertional inactivation of SMc00384*;* Sm$^R$, Neo$^R$ | A. Becker, Germany. |
| SmPI_2011. 123.03G6 | Rm2011 derivative with insertional inactivation of SMc00146; Sm$^R$, Neo$^R$ | This study |

| Plasmids | Relevant characteristics | Source or reference |
|---|---|---|
| pXLGD4 | p*hemA:lacZ* reporter plasmid; Tc$^R$ | 4 |
| pCAP97 | *rhaS* insertion plasmid, p*trp*; Spc$^R$ | 5 |
| pCAP97-SMa2020(WT) | pCAP97 derivative with *sydR* under the control of p*trp*; Spc$^R$ | This study |
| pCAP97-SMa2020*(nodA)* | pCAP97 derivative with *sydR* under the control of p*nodA*; Spc$^R$ | This study |
| pCAP97-SMa2020_C16S | pCAP97 derivative with a *sydR* mutated gene (coding for a SydR protein with a Cys16→Ser exchange) under the control of p*trp*; Spc$^R$ | This study |
| pCAP97-SMa2020_C114S | pCAP97 derivative with a *sydR* mutated gene (coding for a SydR protein with a Cys114→Ser exchange) under the control of p*trp*; Spc$^R$ | This study |
| pCAP97-SMa2020_C16S&C114S | pCAP97 derivative with a *sydR* mutated gene (coding for a SydR protein with Cys16, Cys114 →Ser exchanges) under the control of p*trp*; Spc$^R$ | This study |
| pET28a | Plasmid for overexpression of genes in *E. coli*, generates polyhistidine-tagged fusion proteins; Km$^R$ | Novagen |
| pET28a-SMa2020(WT) | pET28a derivative for overproduction and purification of SydR with an C-terminal polyhistidine tag; Km$^R$ | This study |
| pET28a-SMa2020_C16S | pET28a-SMa2020(WT) derivative coding for a SydR protein with a Cys16→Ser exchange; Km$^R$ | This study |
| pET28a-SMa2020_C114S | pET28a-SMa2020(WT) derivative coding for a SydR protein with a Cys114→Ser exchange; Km$^R$ | This study |
| pET28a-SMa2020_C16S&C114S | pET28a-SMa2020(WT) derivative coding for a SydR protein with Cys16, Cys114 →Ser exchanges; Km$^R$ | This study |
| pJQmp18 | Suicide cloning vector in *S. meliloti* allowing sucrose selection; Gm$^R$ | 6 |
| pJQ'2019-Tc$^R$-2023' | pJQmp18 derivative used for construction of Δ*sydR* mutant; Gm$^R$, Tc$^R$ | This study |
| pTB° | Derivative of the broad-host range cloning vector pBBR1MCS, p*trp*; Spc$^R$, Cm$^R$ | 3 |
| pTBroGFP2-Orp1 | pTB° derivative with *roGFP2-Orp1* under the control of p*trp*; Spc$^R$, Cm$^R$ | 3 |

### REFERENCES


1. Finan TM, Kunkel B, De Vos GF, Signer ER. 1986. Second symbiotic megaplasmid in *Rhizobium meliloti* carrying exopolysaccharide and thiamine synthesis genes. J Bacteriol 167:66–72. https://doi.org/10.1128/jb.167.1.66-72.1986.
2. Pobigaylo N, Wetter D, Szymczak S, Schiller U, Kurtz S, Meyer F, Nattkemper TW, Becker A. 2006. Construction of a large signature-tagged mini-Tn5 tTransposon library and its application to mutagenesis of *Sinorhizobium meliloti*. Appl Environ Microbiol 72:4329–37. https://doi.org/10.1128/AEM.03072-05.
3. Pacoud M, Mandon K, Cazareth J, Pierre O, Frendo P, Alloing G. 2022. Redox-sensitive fluorescent biosensors detect *Sinorhizobium meliloti* intracellular redox changes under free-living and symbiotic lifestyles. Free Rad Biol Med 184:185–95. https://doi.org/10.1016/j.freeradbiomed.2022.03.030.
4. Leong SA, Williams PH, Ditta GS.1985. Analysis of the 5' regulatory region of the gene for delta-aminolevulinic acid synthetase of *Rhizobium meliloti*. Nucleic Acids Res. 13:5965-76. https://doi.doi: 10.1093/nar/13.16.5965.
5. Arango Pinedo C, Gage DJ. 2009. Plasmids that insert into the rhamnose utilization locus, *rha*: a versatile tool for genetic studies in *Sinorhizobium meliloti*. J Mol Microbiol Biotechnol 17(4):201–10. https://doi.org/10.1159/000242446.
6. Quandt J, Hynes M. 1993. Versatile suicide vectors which allow direct selection for gene replacement in gram-bacteria. Gene 127:15–21. https://doi.org/10.1016/0378-1119(93)90611-

**Table S3. Primers used in this study.**

| Primers | 5'-3' nucleotidic sequence |
|---|---|
| PCR primers | |
| *sma2019* up114 | CGGCAAGTCGATAGCGAAG |
| *sma2019* down251 | CCGCCTTCCAGATCAGTACC |
| *sma2019* up172 | GAGTTGTTCGTAGAAGCTGAC |
| *sma2019* down35 | AGATATCTACTCGCAGATTGGC |
| *sma2020_23*F | TCTGGGCTCCTTTGCGC |
| *sma2020_23*R | GGCTTTTTCCTTTCGTTGATAGC |
| *sma2020*+ATG | CACCATATGACAAACCCCATAATG |
| *sma2020* down *Hind*III | TCTAAGCTTCGATCACGGGTAC |
| *sma2020*+SD | TCAGTCGACGGAGGAAACAAAGATGACCAAACCCATAATG |
| *sma2020* down *Kpn*I | TCAGGGTAGAATACACGATC |
| *sma2020*F_Ser46 | GTGTAAATCGCGTAGCTCAACTGATCGTGCAATG |
| *sma2020*R_Ser46 | CATTGCACGATCAGTTGAGCTACGCGATTTACAC |
| *sma2020*F_Ser340 | GTCGCTGAGGCTGCCTGCCCTGTGC |
| *sma2020*R_Ser340 | CGAAGGGCAGGCAGCCTCAGCGAC |
| *sma2023* up144 | GATATCTGGGCTCCTTTGCG |
| *sma2023* down418 | GTCGACGAACTCCGCTGCCG |
| up_*smc03824*F | GTCGACTTTCCTCGTTCTTGATG |
| up_*smc03824*R | GAATTCGAGTTTCCTGGGCAT |
| *smc03824*F | GAGAATTCCTCCGGGACAGTC |
| *smc03825*R | TTCTAGAGCTGCACGATGAAGCAC |
| p*nodA* up+pCAP97 | TCCTCCGTCGACGGTATCGATAAGCTTGTGTACCCGGCAAGTTACAC |
| p*nodA* down+*sma2020* | GGGCTGCAGGAATTCGATATCAAGCTTGGCCCCTAAAACGCATGTG |
| qPCR primers | |
| *sma2023-1*F | CGTCACAATGTCCAGAACC |
| *sma2023-1*R | TCTCGTCTATGTTCGCTCTC |
| *smc00147*F | AAGGTCACGTTGTCCTTTGC |
| *smc00147*R | GATCAGCGTTGCCAGATAGTC |
| *sm00383*F | AAACGCAACGTGCTCAAGTC |
| *sm00383*R | GTGTCGAACACGTCGAAATAGC |
| *smc03825* F2 | CAGTCGATCTTCCTGCTGATG |
| *smc03825* R2 | CCTCATAAGCTCGCCATAGG |
| *smc03979*F | TCGTGGTGGTGGTTATCAAA |
| *smc03979*R | TTCAGACCTTCCCATTGACA |
| *betS* F | TGAGGGAGCAACCAAATACC |
| *betS* R | GCGAAAACCAAGCTACCATC |
| *16S* F | GATAAGCCGAGAGGAAGGTG |
| *16S* R | GTGTAGCCCAGCCCGTAAG |
| *MtENOD11*F | GCCACCATTTCCTAAGCCAC |
| *MtENOD11*R | GGGAAACGAAAACTTGACCC |
| *MtNIN*F | GGAAGATTGAGAGGGGAAGCTT |
| *MtNIN*R | GCAATGTGGGGATTTAGAGATT |
| *MtLYK3*F | CAAGAATTAGCCAAGGCGACA |
| *MtLYK3*R | AGCTCCAAATCCACCTTGACC |
| *MtN20*F | AGATTTCAGCAGATGTGGTAGCGT |
| *MtN20*R | CCCTTCAAACCACCCAAAGAG |
| *MtC27*F | TGAGGGAGCAACCAAATACC |
| *MtC27*R | GCGAAAACCAAGCTACCATC |
| *A38*F | TCGTGGTGGTGGTTATCAAA |
| *A38*R | TTCAGACCTTCCCATTGACA |

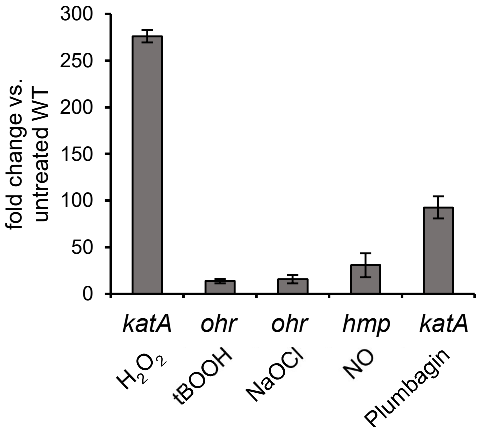


**FIG S1. Determination of oxidant treatments inducing the expression of selected oxidative stress-responsive genes.** RT-qPCR analysis of the expression of *katA*, *ohr*, and *hmp* genes in the WT strain treated with $H_2O_2$ (1 mM) or plumbagin (50 μM) for 10 min, tBOOH (200 μM) or NaOCl (20 μM) for 10 min, and NO (25 μM) for 30 min, respectively. For each condition, transcription levels were normalized to those in untreated WT. The values shown are the means ± SEM of three independent experiments, and significance compared to untreated WT was determined in Student's *t*-test ($p<0.05$).

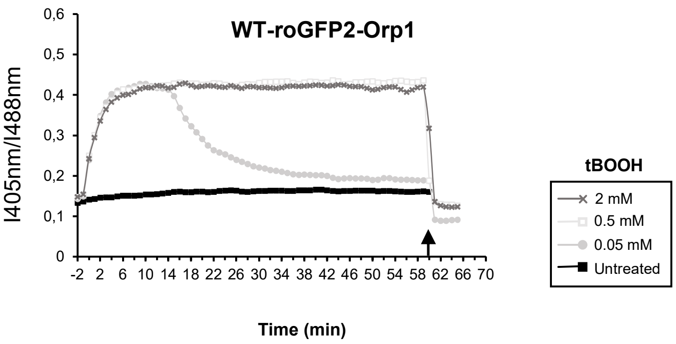


**FIG S2. The oxidation of roGFP2-Orp1 probe by tBOOH is reversible.** Measurements of variations in roGFP2-Orp1 oxidation state were performed in WT treated with various concentrations of tBOOH during 1h, and reversibility of probe oxidation was assessed by adding 10mM of DTT (indicated by the arrow).

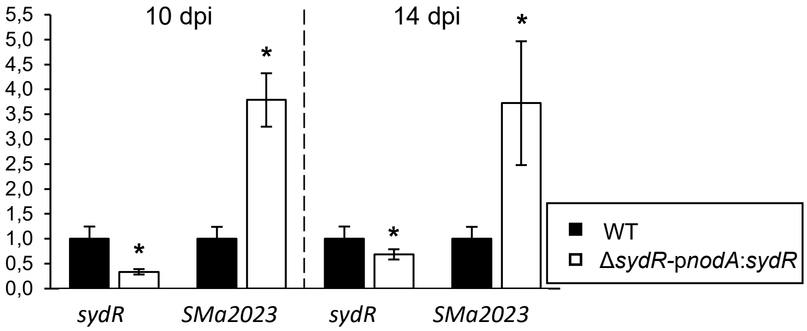


**FIG S3. Validation of the p*nodA*:*sydR* construct.** RT-qPCR analysis of the expression of *sydR* and SMa2023 in nodules infected with WT or Δ*sydR*-p*nodA*:*sydR* strain, at 10 and 14 dpi. For each condition, transcription levels were normalized to those in WT. The values shown are the means ± SEM of three independent experiments, and significance compared to WT was determined in Mann-Whitney test ($p<0.05$).

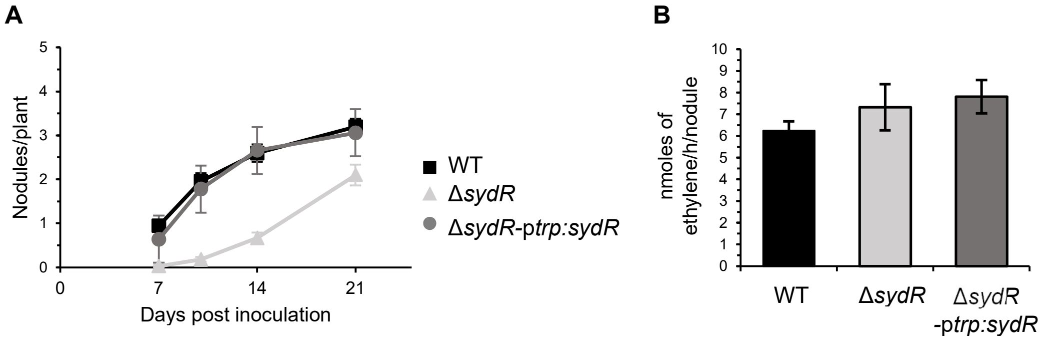


**FIG S4. Δ*sydR* mutant induces the formation of nitrogen-fixing nodules in *M. sativa*.** (A) Nodulation kinetics of *M. sativa* plants inoculated with WT, Δ*sydR* and Δ*sydR*-p*trp:sydR* strains (72 plants in three independent experiments, n=24). The number of nodules per plant is significantly different in roots inoculated with Δ*sydR* as compared to WT at all kinetic points. (B) Nitrogen fixation activity, determined by acetylene reduction assay (ARA) at 21 dpi. The values shown are the means ± SEM of three independent experiments. Nonparametric Kruskal-Wallis and *post-hoc* Conover-Iman tests with Benjamini-Hochberg correction were used to assess the statistical significance of differences compared to the WT strain.

### SUPPLEMENTAL MATERIAL TEXT S1

**Bacterial strains and growth conditions.** *S. meliloti* strains were grown at 30°C in LB medium supplemented with 2.5 mM $MgSO_4$ and 2.5 mM $CaCl_2$ (LBMC), in M9 medium (42.5 mM $Na_2HPO_4$, 22 mM $KH_2PO_4$, 8.5 mM NaCl, 18.8 mM $NH_4Cl$, 1 mM $MgSO_4$, 0.25 mM $CaCl_2$, 1 µg $mL^{-1}$ biotin and 0.2% mannitol) and in M9-CSA (M9 medium supplemented with 0.2% Casamino-Acids). Antibiotics were used at the following concentrations: streptomycin (100 µg $mL^{-1}$), ampicillin (100 µg $mL^{-1}$), kanamycin (25 µg $mL^{-1}$), chloramphenicol (20 µg $mL^{-1}$ for *S. meliloti*, and 10 µg $mL^{-1}$ for *E. coli*), neomycin (200 µg $mL^{-1}$), and spectinomycin (200 µg $mL^{-1}$ for *S. meliloti*, and 50 µg $mL^{-1}$ for *E. coli*).

**Plant growth conditions.** Seeds of *M. truncatula* were scarified with 96% $H_2SO_4$ for 6 min 30 s, and then washed with reverse osmosis water. They were sterilized with 3% NaOCl for 1 min 30 s, washed again in sterile water and left to soak for 3 to 5 h. Finally, they were placed on 0.4% agar plates, synchronized in a cold room at 4°C for 48 h in the dark, and then germinated in a plant culture room. Germinated seeds were grown *in vitro* on Fahraeus medium plates supplemented with 1,5% agar, 0.2 mM $NH_4NO_3$ and 1 mM $CaCl_2$. Seeds of *M. sativa* were sterilized with 3% NaOCl for 10 min, washed in sterile water, placed on 0.4% agar plates and germinated for 48 h. Germinated seeds were thereafter grown in sterilized test tubes containing Fahraeus *slant agar.*

For *M. truncatula* or *M. sativa* inoculation, bacterial cultures grown to an $OD_{600}$ of 0.8-1 in LBMC with appropriate antibiotic(s) were washed twice in sterile water and resuspended at an $OD_{600}$ of 0.01 or 0.2. Four-days-old (*M. truncatula*) or two-days-old (*M. sativa*) seedlings were inoculated with 200 µL of bacterial suspension per plant.

All plants were grown under a 16 h light (23°C) - 8 h dark (20°C) photoperiod.

**Construction of *S. meliloti* mutants.** To generate in-frame 398 bp deletion in the SMc03824 ORF, DNA fragments encompassing the 5' and 3' ends of SMc03824 were first amplified from *S. meliloti* genomic DNA with two pairs of primers, up_*smc03824*F/up_*smc03824*R, and *smc03824*F/*smc03825*R, respectively (Table S2). Each fragment was thereafter cloned into pJET2.1, ligated together, and inserted as a *Sal*I/*Xba*I fragment into the suicide vector pJQmp18 (1). The recombinant plasmid was introduced into the Rm2011 recipient strain by conjugation with the helper strain MT616, as previously described (2). The double-recombinant clones were selected as $sucrose^R$ on M9 minimal medium.

The SMc00146 insertion mutant was obtained by conjugation with SmPI_S17-1.123.03G6, which carries an internal fragment of the SMc00146 ORF, and was kindly provided by Anke Becker (https://www.cebitec.uni-bielefeld.de/CeBiTec/rhizogate).

The Δ*sydR* mutant is a deletion-insertion mutant, in which the *sydR* ORF was deleted and replaced by a tetracycline resistance ($Tc^R$) cartridge. Firstly, the upstream and downstream flanking regions of *sydR* were amplified with specific primer pairs (*sma2019* up172/*sma2019* down35, and *sma2023* up144/*sma2023* down418, Table 2). Secondly, the fragment amplified with *sma2019* up172/*sma2019* down35 was inserted as an *Eco*RV fragment into the *Sma*I site of pJQmp18, giving pJQ'2019. In parallel, the fragment amplified with *sma2023* up144/*sma2023* down418 was cloned into pJET2.1, and the $Tc^R$ cartridge from pHP45-Ω-Tc (3) was inserted into the *EcoR*V site located at the 5' end of SMa2023 sequence. Finally, the *Bgl*II-*Xba*I fragment carrying $Tc^R$-SMa2023 was subcloned between the *Bam*HI-*Xba*I sites of pJQ'2019, giving pJQ'2019-$Tc^R$-2023'. This plasmid was conjugated into the Rm2011 by triparental conjugation as above, and the double-recombinants were selected as $Tc^R$, $sucrose^R$.

**Molecular cloning and mutagenesis of *sydR*.** For the construction of Δ*sydR* strains expressing *sydR* constitutively, wild-type and mutated *sydR* genes cloned in pJET1.2/blunt were subcloned into the *Kpn*I and *Sal*I sites of pCAP97, downstream of the *Salmonella typhimurium trp* promoter (4, 5). The resulting pCAP97-SMa2020(WT), pCAP97-SMa2020_C16S, pCAP97-SMa2020_C114S, and pCAP97-SMa2020_C16S&C114S were conjugated into the Δ*sydR* mutant strain. The pCAP97 contains a *rhaS* fragment of the *S. meliloti* rhamnose locus, promoting its integration by homologous recombination into the chromosome. The proper integration of plasmids in the ex-conjugant genomes was confirmed by testing the loss of ability to use rhamnose as a carbon source and by colony PCR.

To construct the Δ*sydR* strain expressing SMa2020 from the p*nodA* promoter, the pCAP97-p*trp*-SMa2020 was converted into pCAP97-p*nodA*-SMa2020 using T5-Exonuclease Dependent DNA Assembly (TEDA) as described by Xia *et al*, (6). The p*trp* promoter was first removed from the pCAP97-SMa2020 by *Hind*III restriction. The p*nodA* was amplified by PCR using *S. meliloti* genomic DNA as a template, p*nodA* up+pCAP97/p*nodA* down+*sma2020* primers and Phusion DNA polymerase (New England Biolabs). The linearized plasmid and the PCR fragment carrying the p*nodA* promoter were added to 5X TEDA reaction solution (New England Biolabs), with a molar ratio vector vs insert of 1:4. The resulting pCAP97-p*nodA*-SMa2020 was then recombined into the Δ*sydR* genome as described above.
All recombination events were controlled by PCR colony.

To produce recombinant SydR wild-type and variant proteins, the coding region of the *sydR* gene was first amplified by PCR using Rm2011 genomic DNA as template and primer pairs *sma2020*+SD/*sma2020* down *Kpn*I or *sma2020*+ATG /*sma2020* down *Hind*III. Both PCR fragments were directly cloned into pJET1.2/blunt cloning vector (ThermoScientific), generating pJET1.2-SMa2020(WT) and pJET1.2-SMa2020(ATG), respectively. pJET1.2-SMa2020(WT) was used as a template for *sydR* mutagenesis and subcloning into the suicide vector pCAP97. pJET1.2-SMa2020(ATG) was digested with *Nde*I and *Xho*I and the *sydR* fragment ligated into the *Nde*I-*Sal*I sites of the pET28a vector, giving pET28a-SMa2020(WT) which provides a 6 histidine-Tag at the C-terminus of the expressed proteins.

To perform the *sydR* mutagenesis, the Quick Change II site-directed mutagenesis kit (Stratagene) was used to replace one or both cysteine residues of SydR (C16 and C114) with serine. The plasmid pET28a-SMa2020 or pJET1.2-SMa2020 was used as a PCR template with the corresponding complementary mutagenic primers: *sma2020F_Ser46*/*sma2020R_Ser46* and *sma2020F_Ser340*/*sma2020R_Ser340*, resulting in four vectors with the corresponding mutated SMa2020 genes, pET28a-SMa2020_C16S, pET28a-SMa2020_C114S, pJET1.2-SMa2020_C16S, and pJET1.2-SMa2020_C114S. To produce the SydR mutant protein with mutation in both residues, the complementary mutagenic primers *sma2020F_Ser340* /*sma2020R_Ser340* were used, but the plasmid either pET28a-SMa2020_C16S or pJET1.2-SMa2020_C16S was used as a PCR template. All mutations generating single or double aminoacid substitutions were verified by DNA sequencing.

**RNA extraction and RT-qPCR assays.** Total RNAs from frozen bacterial pellets were extracted using the RNeasy kit (Qiagen, USA), then treated with DNase I to remove potential contaminating DNA. In order to quantify gene expression *in planta*, freshly harvested roots and nodules were frozen in liquid nitrogen and ground to a fine powder, and total RNA was extracted from 100 mg of powder using RNAzol®RT (Molecular Research Centre, Inc., Cincinnati, USA) according to the manufacturer's instructions.

The cDNAs were generated using the GoScript Reverse Transcription kit (Promega, USA). The synthesized cDNA was diluted 20- to 40-fold and used as template to analyze relative gene expression. The quantitative PCR was performed with GoTaqR qPCR Master Mix kit and gene specific primer pairs (Table S3) using the Agilent AriaMX thermal cycler. The qPCR program consisted of an initial denaturation at 95°C for 3 min, followed by 40 cycles of 3 sec at 95°C and 30 sec at 60°C, and melting curves from 65°C to 95°C in increments of 0.5°C. Bacterial reference genes (16S rRNA, SMb20333 (*betS*), and SMc03979 or plant reference genes (*A38* and *MtC27*; 7) were used to normalize the data. All measurements were performed in biological and technical triplicates. The expression fold change was calculated using the $2^{-\Delta\Delta Ct}$ method (8). Primers used in RT-qPCR are listed in Table S4.

**Purification of SydR wild-type and mutant proteins.** For recombinant protein production, BL21 (DE3) pLysS competent cells were transformed with the pET28a-SMa2020, pET28a-SMa2020_C16S, pET28a-SMa2020_C114S, and pET28a-SMa2020_C16S&C114S plasmids. To produce the recombinant proteins, overnight bacterial cultures were subcultured into 100 ml of fresh medium and incubated at 37°C until $OD_{600}$ of ~0.5. Afterward, the cultures were induced with 1mM isopropyl-β-D-thiogalactopyranoside (IPTG), incubated for 3h at 37°C with vigorous shaking, and then harvested by centrifugation. The pellets were resuspended in lysis buffer (50 mM $NaH_2PO_4$ pH 8, 300 mM NaCl) and lysed by sonication (at 60% amplitude for 2 min at regular intervals of 15 seconds). The recombinant proteins SydR' were purified from the soluble fraction by HisPur™ Ni-NTA resin affinity chromatography according to the manufacturer's recommendation (Thermoscientific, USA). The Amicon ultra centrifugal desalting columns with 10 nominal molecular weight limit (NMWL) were used to exchange buffers and make the purified proteins appropriate for subsequent use. The purity of proteins was analyzed by 15% sodium dodecyl sulphate polyacrylamide gel electrophoresis (SDS-PAGE), and their concentrations were measured by the Bradford method (Biorad reagent).

**Microscopy and histology analysis.** For GUS staining, roots were were fixed with 90% acetone for 1h at −20°C, rinsed three times in 100 mM phosphate buffer (pH 7.4), and incubated overnight at 37°C in GUS staining solution (phosphate buffer supplemented with 0.05% 5-bromo-4-chloro-3-indolyl-β-D-glucuronic acid (X-gluc), 0.5 mM $K_3Fe(CN)_6$ (ferricyanide), and 0.5 mM $K_4Fe(CN)_6$ [ferrocyanide]), as previously described. Roots were washed three times in phosphate buffer, cleared by a 3 min treatment in 3.2% NaOCl, and washed again in phosphate buffer, before observation under a transmission light microscope (Zeiss Axioplan II).

Roots inoculated with bacterial strains expressing the p*hemA:lacZ* fusion were harvested at 4 and 10 dpi (n=6 roots per time point). They were fixed with 2% glutaraldehyde for 1.5 hours under vacuum, rinsed three times in 100 mM phosphate buffer (pH 7.4) and incubated in β-gal staining solution (0.04% 5-bromo-4-chloro-3-indolyl-β-D-galactopyranoside (X-gal), 2.5 mM $K_3Fe(CN)_6$, 2.5 mM $K_4Fe(CN)_6$ in phosphate buffer) for 2 h at 30°C. As above, roots were washed before observation under a transmission light microscope. Nodules (14 dpi) were harvested and embedded in 6% agar solution, then 100 µm (nodules from the WT strain) and 120 µm (nodules from the mutant strain) sections were made with a vibratome (Leica VT1200S) sectioning. Nodule sections were submitted to the same staining protocol as described for roots.

### REFERENCES

1. Quandt J, Hynes M. 1993. Versatile suicide vectors which allow direct selection for gene replacement in gram- bacteria. Gene 127:15–21. https://doi.org/10.1016/0378-1119(93)90611-6.

2. Finan TM, Kunkel B, De Vos GF, Signer ER. 1986. Second symbiotic megaplasmid in *Rhizobium meliloti* carrying exopolysaccharide and thiamine synthesis genes. J Bacteriol 167:66–72. https://doi.org/10.1128/jb.167.1.66-72.1986.
3. Fellay R, Frey J, Krisch H. (1987). Interposon mutagenesis of soil and water bacteria: a family of DNA fragments designed for *in vitro* insertional mutagenesis of gram-negative bacteria. Gene 52:147–154. https://doi.org/10.1016/0378-1119(87)90041-2.
4. Arango Pinedo C, Gage DJ. 2009. Plasmids that insert into the rhamnose utilization locus, *rha*: a versatile tool for genetic studies in *Sinorhizobium meliloti*. J Mol Microbiol Biotechnol 17:201–210. https://doi.org/10.1159/000242446.
5. Gage, Daniel J., Tanya Bobo, Sharon R. Long. 1996. Use of green fluorescent protein to visualize the early events of symbiosis between *Rhizobium meliloti* and alfalfa (*Medicago sativa*). J Bacteriol 178:7159–7166. https://doi.org/10.1128/jb.178.24.7159-7166.1996.
6. Xia Y, Li K, Li J, Wang T, Gu L, Xun L. 2019. T5 exonuclease-dependent assembly offers a low-cost method for efficient cloning and site-directed mutagenesis. Nucleic Acids Res 47(3). https://doi.org/10.1093/nar/gky1169.
7. Del Giudice J, Cam Y, Damiani I, Fung-Chat F, Meilhoc E, Bruand C, Brouquisse R, Puppo A, Boscari A. 2011. Nitric oxide is required for an optimal establishment of the *Medicago truncatula-Sinorhizobium meliloti* symbiosis. New Phytol 191:405-417. https://doi.org/10.1111/j.1469-8137.2011.03693.x
8. Livak K, Schmittgen T. 2001. Analysis of relative gene expression data using real-time quantitative PCR and the $2^{-\Delta\Delta Ct}$ method. Methods 25 :402-408. https://doi.org/10.1006/meth.2001.1262.
